# Discovery of Novel Inhibitors of HMG-CoA Reductase using Bioactive Compounds isolated from Cochlospermum Species through Computational Methods

**DOI:** 10.1101/2025.01.19.633828

**Authors:** Toba Isaac Olatoye

## Abstract

Cholesterol biosynthesis is a critical pathway in cellular metabolism, with 3-hydroxy-3-methylglutaryl coenzyme-A reductase (HMGR) catalyzing its committed step. The inhibition of HMGR has been widely explored as a therapeutic target for managing hypercholesterolemia, and statins are the most commonly used competitive inhibitors. However, the search for novel, natural inhibitors of HMGR is still a vital area of research, especially in light of the adverse effects of the prolonged use of statins. In this study, the potential of 84 phytochemicals isolated from two species of *Cochlospermum* (C. *planchonii* and C. *tinctorium*) reported in the literature, was investigated as novel inhibitors of human HMGR using molecular docking techniques. The phytochemicals were screened for their drug-likeness and ADMET properties in accordance with Lipinski’s rule of five, and 32 of them were docked against the HMG-binding portion of the enzyme’s active site, together with its native ligand and 6 known statins serving as control ligands. The docking results revealed that 10 of the compounds exhibited strong binding affinities and interactions with the HMG-binding pocket of the enzyme, comparable to or exceeding those of the control ligands. This evidence strongly suggests their potential as effective inhibitors of HMGR. For the first time, the findings from this research identify and directly implicate the specific bioactive compounds of C. *planchonii* and C. *tinctorium* capable of exerting cholesterol-lowering effects in humans, while validating previously reported studies on the efficacy of these plants’ extracts used in West African traditional medicine to manage dyslipidemia, among a host of other ailments. The compounds identified may serve as promising drug candidates which can be further optimized and developed as novel, natural inhibitors of HMGR.

## Introduction

Cholesterol is a vital component of cellular membranes and serves as a precursor for the biosynthesis of steroid hormones, bile acids, and vitamin D. However, elevated levels of cholesterol, especially low-density lipoprotein (LDL) cholesterol, are strongly associated with the development of atherosclerosis and cardiovascular diseases (CVDs), which are leading causes of morbidity and mortality worldwide (Goldstein and Brown, 2015). Although lifestyle changes in individuals, such as exercise, healthy diets, and drug therapies-particularly statins, have been touted as effective in the prevention and management of hypercholesterolemia and its attendant cardiovascular complications (Goldstein and Brown, 2015; Ference *et al*., 2017), the challenge is yet far from over. These conditions still remain major global concerns, particularly in developed countries like the United States, where about 48% of adults are currently affected (Virani *et al*., 2023).

HMG-CoA (3-hydroxy-3-methylglutaryl-coenzyme A) reductase (HMGR), the rate-limiting enzyme in the mevalonate pathway, catalyzes the four-electron reductive deacylation of HMG-CoA to mevalonate, which is a crucial precursor of cholesterol biosynthesis in human (Istvan *et al*, 2000). Statins, a class of synthetic drugs with inhibition constant (Ki) values in the nanomolar range, are competitive inhibitors of HMGR widely used to lower serum cholesterol levels in human (Istvan and Deisenhofer, 2001). These drugs occupy the catalytic portion of the enzyme where the substrate, HMG-CoA binds, thus blocking its access to the active site (Fig.1). Near the carboxyl terminus of human HMGR, several catalytically relevant amino acid residues representing the HMG-binding pocket are disordered in the enzyme-statin complex. If these residues were not flexible, they would sterically hinder the binding of statins (Istvan and Deisenhofer, 2001). All statins share an HMG-like moiety, with rigid, hydrophobic groups that are covalently linked to them and may be present in inactive lactone form. In vivo, these drugs are enzymatically hydrolyzed to their active hydroxy-acid forms (Corsini *et al*., 1995). In addition to lowering cholesterol, statins appear to have other functions, including the nitric oxide-mediated promotion of the growth of new blood vessels (Kureishi *et al*., 2000), stimulation of bone formation (Mundy *et al*., 1999), protection against oxidative modification of low-density lipoprotein, and anti-inflammatory effects with a reduction in C-reactive protein levels (Davignon and Laaksonen, 1999). Nevertheless, the use of statins is often limited by their side effects such as myopathy, liver and kidney dysfunction, and an increased risk of diabetes (Tailor *et al*., 2013; Collins *et al*., 2016; Cho *et al*., 2015).). These limitations have necessitated the search for alternative cholesterol-lowering agents, especially those from natural sources, which may offer safer and more effective therapeutic needs.

**Figure 1:**
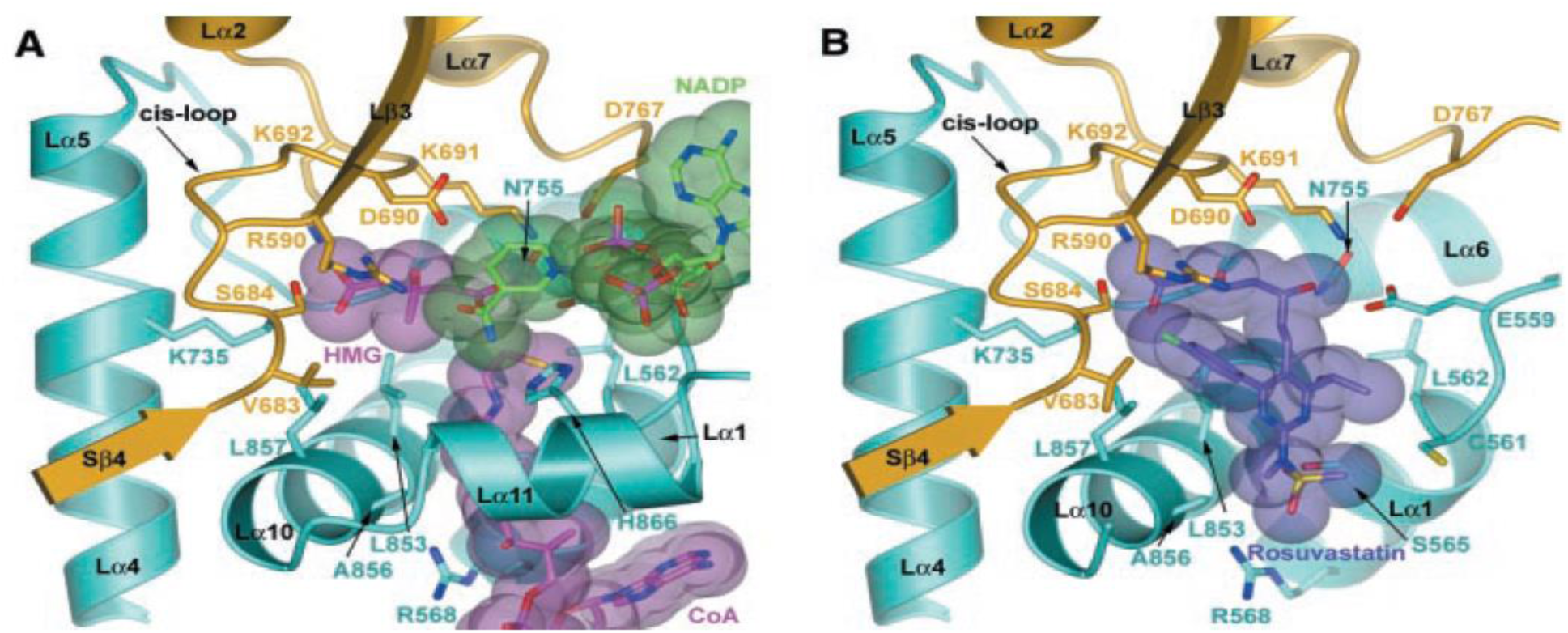
**(A)** Active site of human HMGR in complex with HMG, CoA, and NADP^+^. **(B)** Binding of rosuvastatin to HMGR. The active site is located at a monomer-monomer interface. One monomer is colored yellow, the other monomer is in blue. Selected side chains of residues that contact the substrates or the statin are shown in a ball-and-stick representation. Secondary structure elements are marked by black labels. HMG and CoA are colored in magenta; NADP is colored in green. Rosuvastatin is colored in purple. Statins exploit the conformational flexibility of HMGR to create a hydrophobic binding pocket near the active site. Single-letter abbreviations for the key amino acid residues forming the binding pocket of HMG (where rosuvastatin binds colored yellow) are as follows: K, Lys; D, Asp; R, Arg; S, Ser; V, Val (Istvan and Deisenhofer, 2001).

The three-dimensional (3D) crystal structure of human HMGR (PDB ID: 1HWK), is a tetramer (subunits A: PRO442–HIS861; B: SER463–GLY860; C: LEU462–GLY860; D: SER463–GLY860) that contains the catalytic domains of HMGR in complex with four atorvastatin molecules at the interface of two adjacent monomers (https://www.rcsb.org/structure/1HWK; Istvan and Deisenhofer, 2001). Structurally, these domains are divided into three sub-domains namely: an ‘N-domain’ (residues 460-567), which connects the catalytic portion of the enzyme to the membrane, a large ‘L-domain’ (residues 528–590 and 694–872), and a small ‘S-domain’ (residues 592-682). The amino acid residues of the L- and S-domains form the two active sites; the HMG-binding pocket characterized by a narrow cis-loop (residues 682**-**694) formed between the S- and L-domain, and the NADPH-binding site containing residues 592-682 of the S-domain which are also inserted into the L-domain (Istvan and Deisenhofer, 2001; Istvan *et al*., 2000) (Fig. 1 and 2). Since all statins share HMG-like moieties which enable them to compete with HMG-CoA by sterically preventing its binding at the cis-loop, it is imperative to computationally explore this mode and mechanism of inhibition to determine whether the phytochemicals of interest may exhibit similar interactions at this narrow binding site.

**Figure 2:**
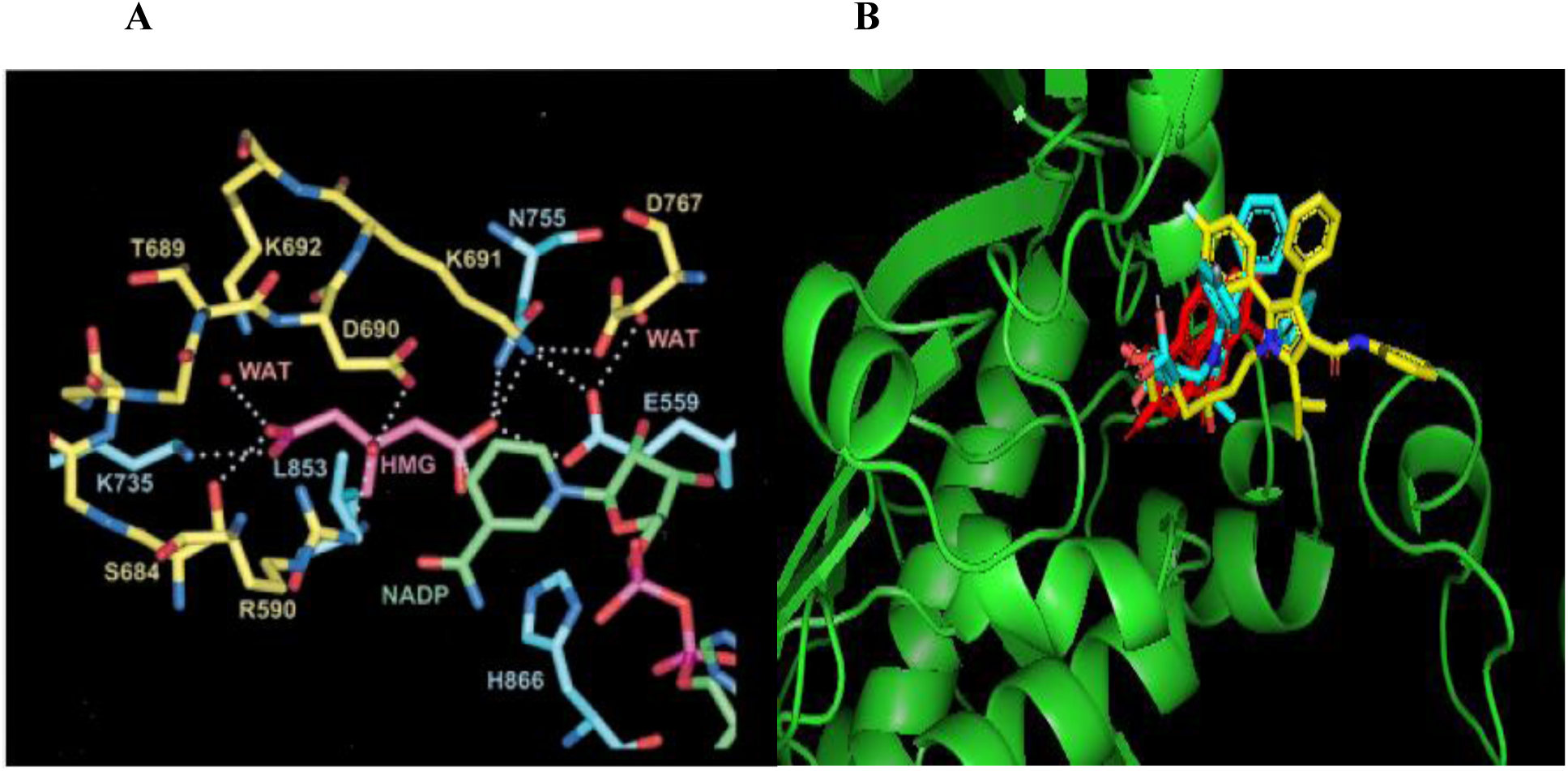
**(A)** HMG-binding site of human HMGR. The stereo diagram shows single letter-abbreviations of residues involved in HMG-binding based on the crystal structure of HMGR (PDB ID: 1HWK) at a resolution of 2.0 Å. Residues from monomer α are in yellow and residues from monomer β are in blue. HMG is magenta and NADP^+^ is in green. All distances within 3.0 Å or closer are indicated by dotted lines (Istvan *et al*., 2000). **(B)** 3D representation showing the binding modes of co-crystallized atorvastatin (yellow), cerivastatin-ID_446156 (cyan), and hit compound, 3-O-methylellagic acid-ID_13915428 (red), at the HMG-binding site of HMGR.

*C. planchonii* and *C. tinctorium*, two species of *Cochlospermum*, are plants extensively used in West African herbal medicine to manage several ailments (Lamien-Meda *et al*., 2015; Dall’Acqua, *et al*., 2020; Ahmad *et al*., 2021; Haidara *et al*., 2016; Johnson-Fulton and Watson, 2018). They are known for their rich phytochemical constituents like, tannins, saponins, carotenoids, triterpenoids, flavonoids, and other polyphenolic compounds, which exhibit various pharmacological activities including antimalarial, antidiabetic, antioxidant, anti-inflammatory, antimicrobial, and enzyme-inhibitory properties (Dall’Acqua, *et al*., 2020; Ballin *et al*., 2002; Habtemariam, 2011; Benoit-Vical *et al*., 2001; Tijjani *et al*., 2009; Etuk *et al*., 2009; Nergard *et al*., 2005; Musa, 2012; Nafiu et al., 2011). Other studies have also demonstrated the antihyperlipidemic and cholesterol-lowering potential of their extracts (root, rhizomes, and leaf) (Nafiu *et al*; 2020; Ahmad *et al*., 2021), suggesting they contain bioactive compounds capable of managing lipid disorders. As far as the literature is concerned, no specific compounds isolated from *C. planchonii* and *C. tinctorium* have been directly studied or linked as potential inhibitors of HMGR. However, their phytochemicals, mostly reported in the literature to possess antioxidant, antimicrobial, anti-inflammatory, and enzyme-inhibitory activities, are thought to be the significant contributors to the plants’ lipid-lowering ability (Nafiu *et al*; 2020; Ahmad *et al*., 2021; Da-Costa-Rocha *et al*., 2014; Dall’Acqua *et al*., 2020).

To explore the efficacy of these phytochemicals as potential inhibitors of human HMGR, molecular docking tools were utilized in this study. Molecular docking is a computational technique used to predict the preferred orientation of a molecule (ligand) when it binds to a target protein, allowing researchers to assess the binding mode and affinity, as well as the chemical interactions between the ligand and the enzyme’s active site as they form a complex (Morris *et al*., 2009). Adopting this approach helps in evaluating the behavior and mechanism of catalysis, stability, flexibility, and conformational changes of the phytochemical-HMGR complexes in a context that is relevant to HMGR inhibition.

## Materials and Methods

### Phytochemical Selection

The selection of phytochemicals for this study was guided by a comprehensive review of existing literature on high-performance liquid chromatography (HPLC) and gas chromatography-mass spectrometry (GC-MS) analyses of various extracts from *C. planchonii* and *C. tinctorium*. This search focused on peer-reviewed articles on both the phytochemical constituents of these plants and their pharmacological properties (Lamien-Meda *et al*., 2015; Dall’Acqua *et al*., 2020; Yahaya *et al*., 2020). The 84 phytochemicals (32 from *C. planchonii*, Table 1; 52 from *C. tinctorium*, Table 2) used for this computational study were selected based on their reported bioactivities, structural integrity, molecular weight, and availability of data regarding their molecular structures. Their two-dimensional (2D) structures were retrieved in structure data file (SDF) format 5^th^ August, 2024 from the PubChem database (https://pubchem.ncbi.nlm.nih.gov) and concatenated using OpenBabel software (https://openbabel.org), before being used for molecular docking studies.

**Table 1:**
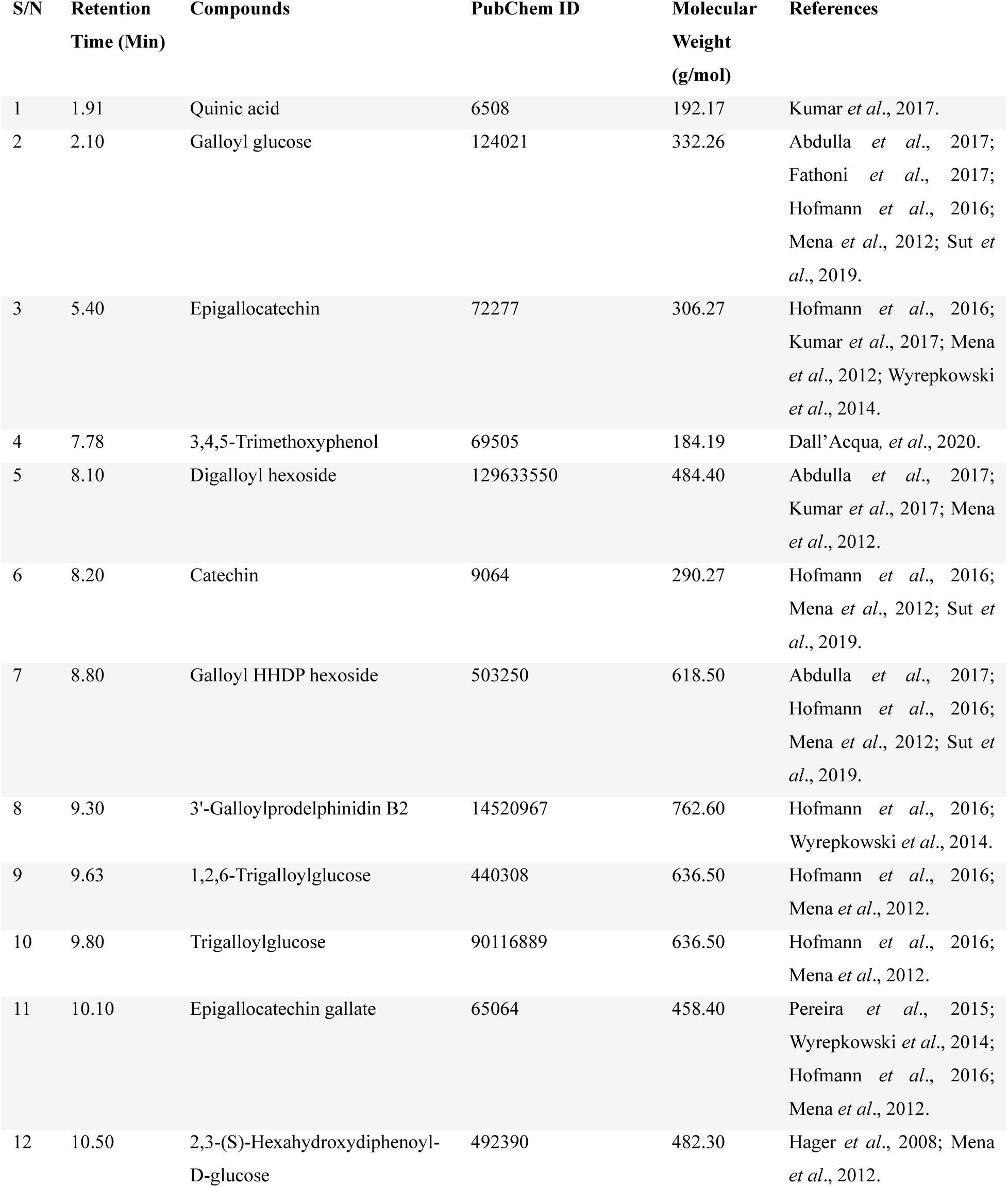
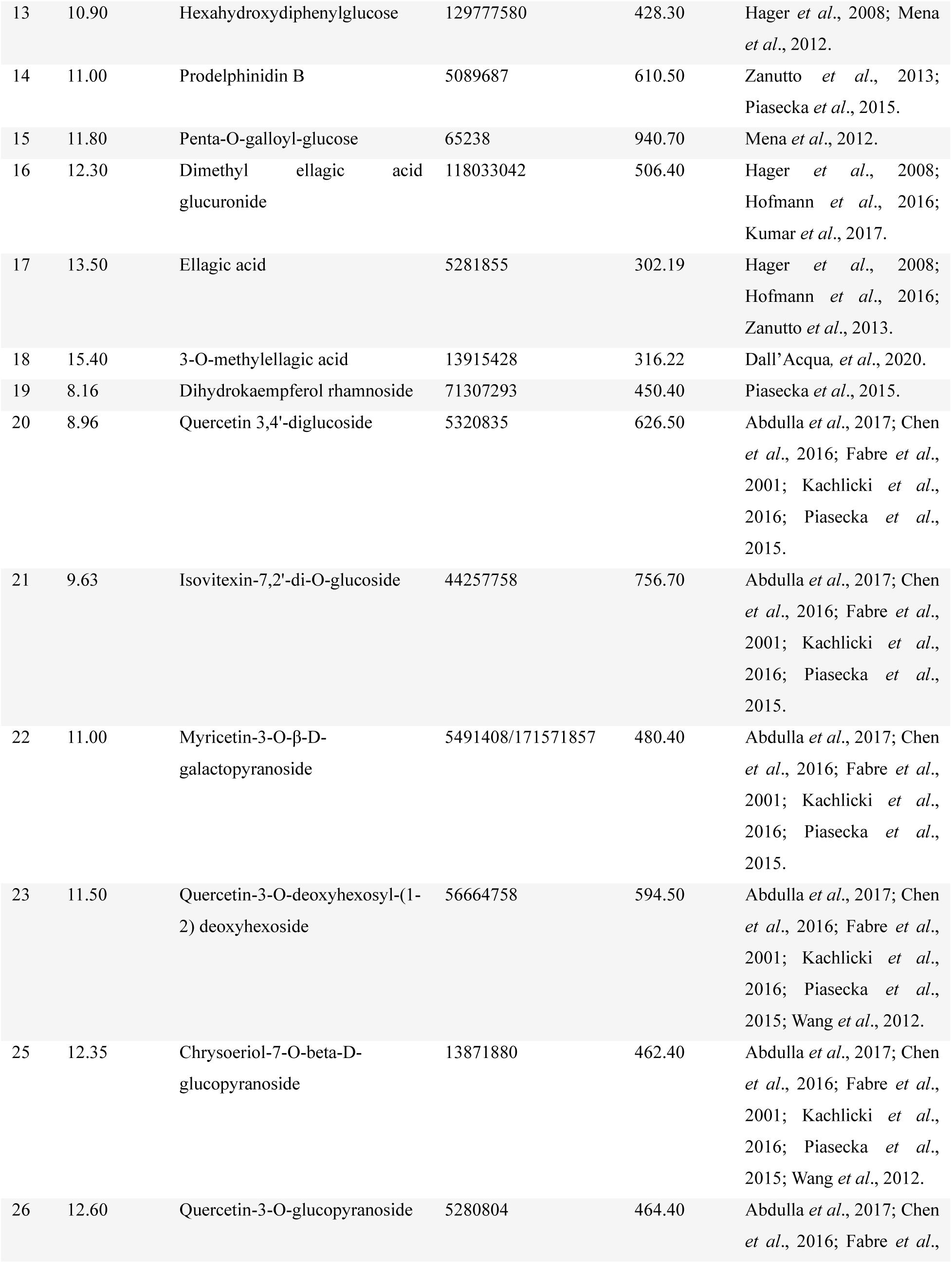
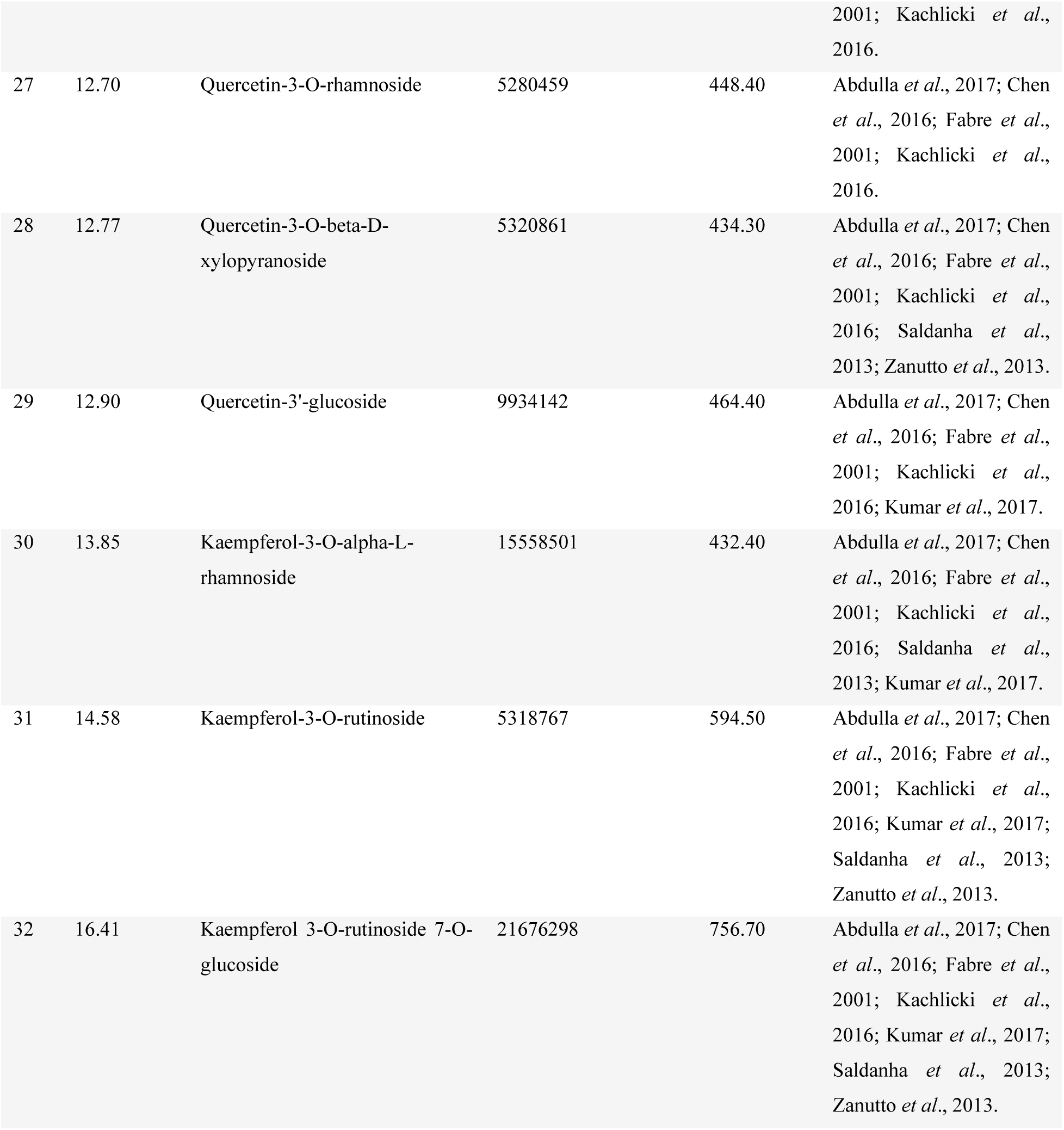
Phytochemicals identified from HPLC analysis of *C. planchonii* using different extraction methods: Maceration (MAC); Homogenization (HAE); Sonification (SON); Soxhlet (SOX)

**Table 2a:**
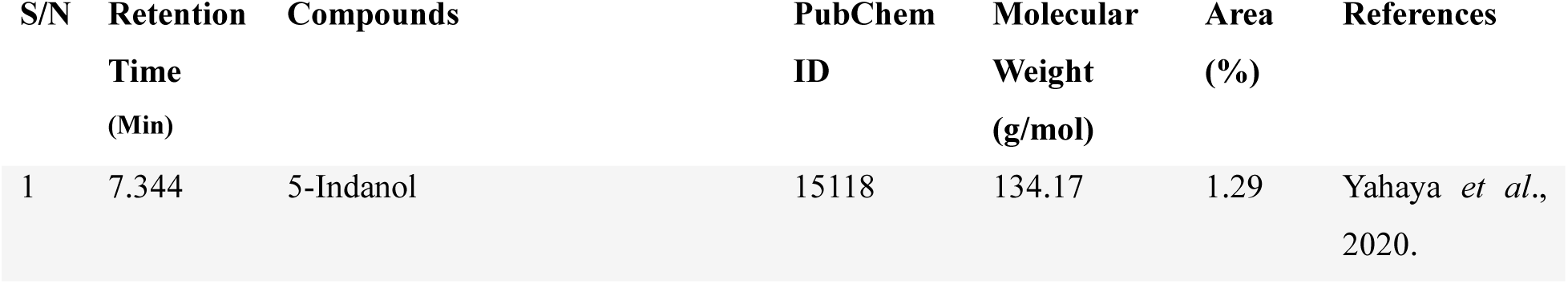
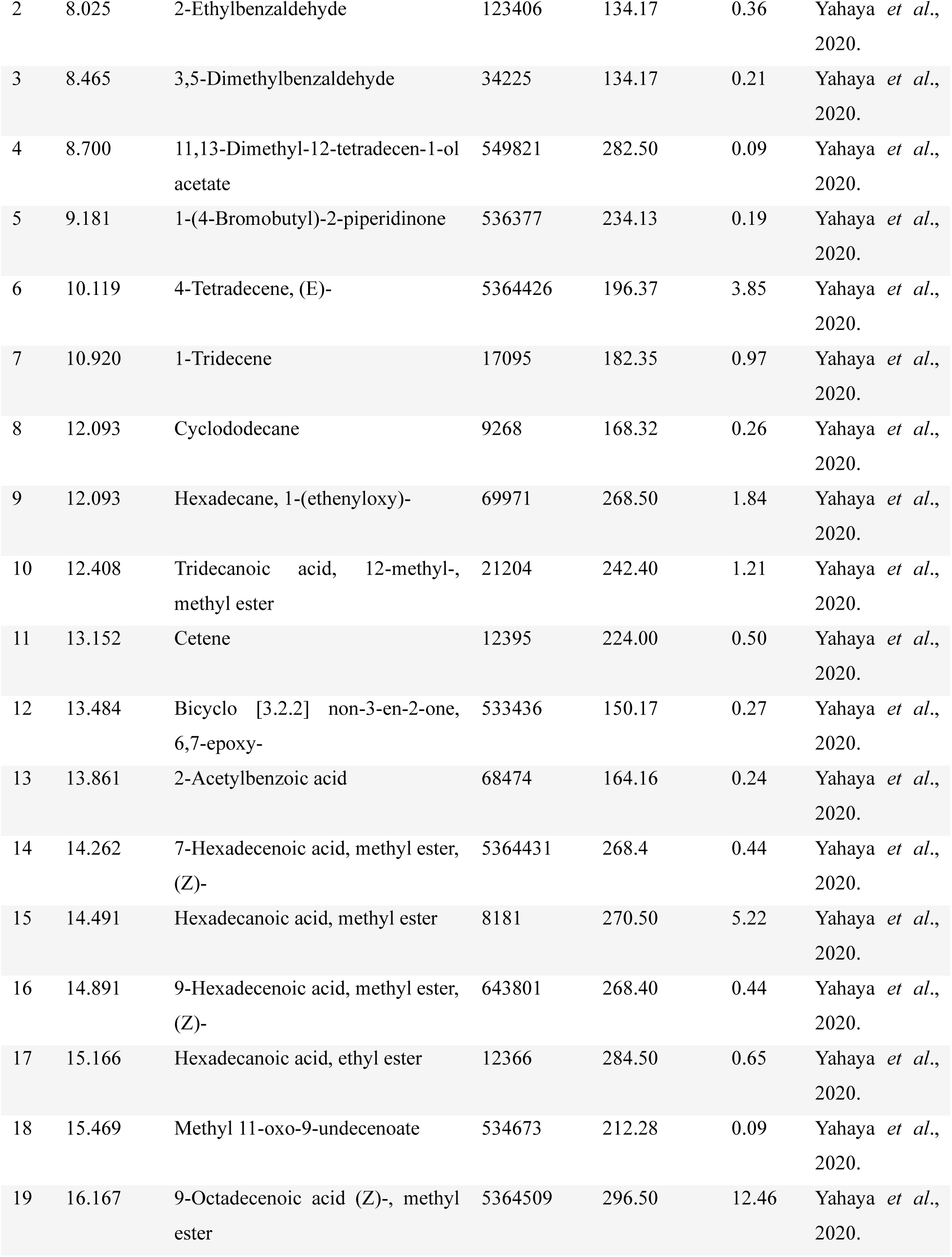
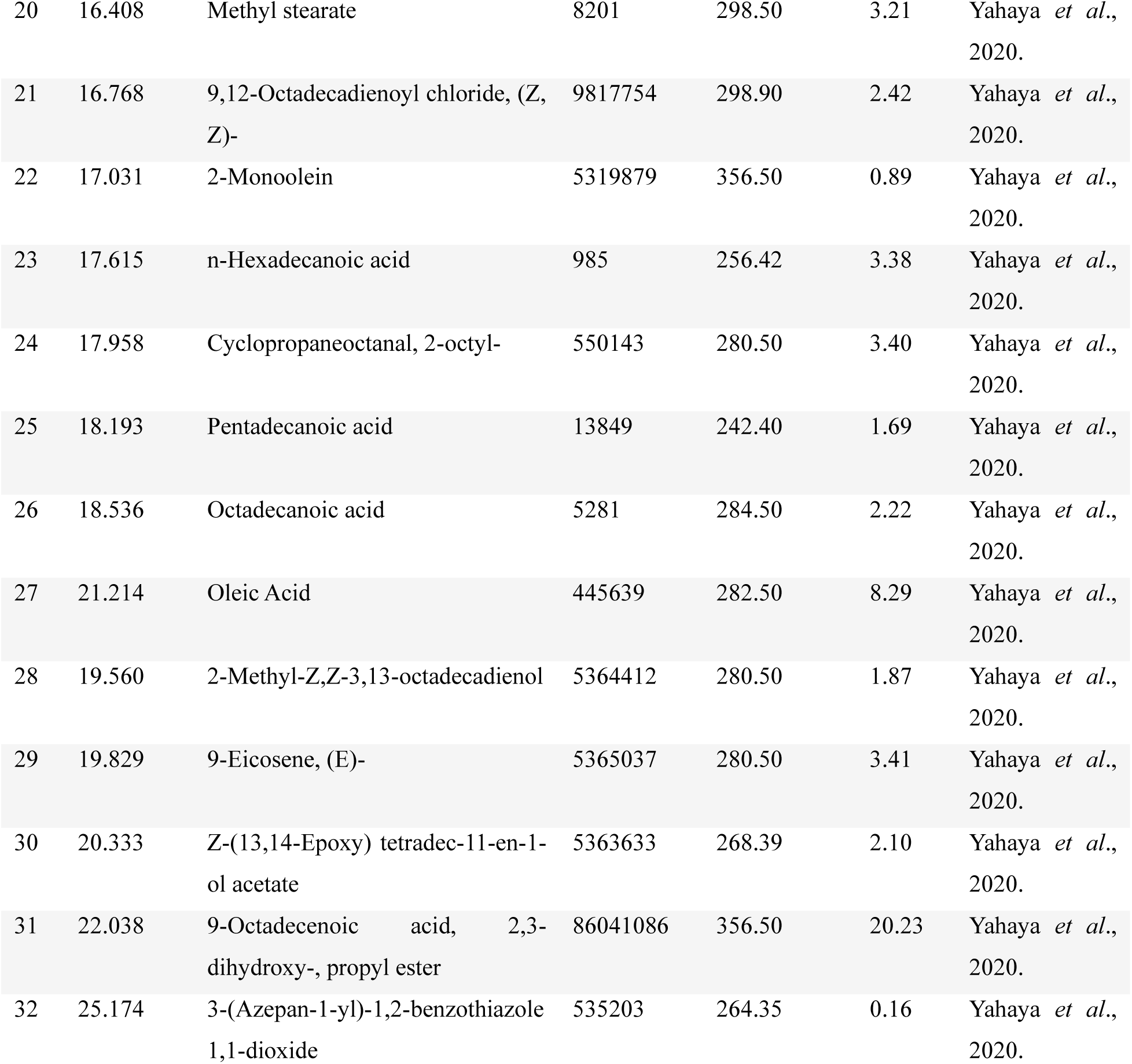
GC-MS analysis of phytochemicals identified in the ethanolic extract of *C. tinctorium*.

**Table 2b:** **HPLC-UV analysis of phytochemicals identified in the ethanolic/methanolic extract of *C. tinctorium***

| S/N | Compounds | PubChem ID | Molecular Weight (g/mol) | References |
| --- | --- | --- | --- | --- |
| 1 | 3-O-p-trans-Coumaroylalphitollic acid | 131751948 | 618.80 | Akinloye *et al.*, 2012; Ballin *et al.*, 2002. |
| 2 | Alphitolic acid | 12305768 | 472.70 | Akinloye *et al.*, 2012; Ballin *et al.*, 2002. |
| 3 | Betulinic acid | 64971 | 456.70 | Akinloye *et al.*, 2012; Ballin *et al.*, 2002. |
| 4 | 3-tetradecanone | 69418 | 212.37 | Benoit-Vical *et al.*, 2001. |
| 5 | 3-hexadecanone | 87789 | 240.42 | Benoit-Vical *et al.*, 2001; Diallo *et al.*, 1991. |
| 6 | Arjunolic acid | 73641 | 488.70 | Diallo *et al.*, 1989. |
| 7 | Carotenoid | 152743345 | 568.90 | Diallo *et al.*, 1991. |
| 8 | 3,7-Di-O-Methylquercetin | 5280417 | 330.29 | Diallo *et al.*, 1992. |
| 9 | 3,4'-O-Dimethylquercetin | 44446550 | 332.30 | Diallo *et al.*, 1992. |
| 10 | Cochloxanthin<br>(2E,4E,6E,8E,10E,12E)-13-[(1S)-1-hydroxy-2,6,6-trimethyl-4-oxocyclohex-2-en-1-yl]-2,6,11-trimethyltrideca-2,4,6,8,10,12-hexaenoic acid | 101202074 | 396.50 | Diallo *et al.*, 1992; Ballin *et al.*, 2002. |
| 11 | 2-Tridecanone | 11622 | 198.34 | Diallo *et al.*, 1991. |
| 12 | 1-Dodecanol | 8193 | 186.33 | Diallo *et al.*, 1991. |
| 13 | 1-Tetradecanol | 8209 | 214.39 | Diallo *et al.*, 1991. |
| 14 | 2-Pentadecanone | 61303 | 226.40 | Diallo *et al.*, 1991. |
| 15 | 2-Stearoylglycerol | 79075 | 385.80 | Diallo *et al.*, 1991. |
| 16 | 2-Octadecanone | 162822 | 284.50 | Diallo *et al.*, 1991. |
| 17 | 3-Octadecanone | 87532 | 268.50 | Diallo *et al.*, 1991. |
| 18 | 1-Hydroxy-3-hexadecanone | 15726205 | 256.42 | Diallo *et al.*, 1991. |
| 19 | 1-Nonadecanol | 80281 | 284.50 | Diallo *et al.*, 1991. |
| 20 | 1-Hydroxy-1-phenyl-3-octadecanone | 131751844 | 360.60 | Diallo *et al.*, 1991. |

### Virtual Screening

DataWarrior (https://www.openmolecules.org/datawarrior) is excellent for managing and screening large libraries of compounds based on their chemical properties (Sander *et al*., 2015). The software was used to narrow down the large pool of 84 potential drug candidates, ensuring that only the most promising ones make it to the docking step. This approach helps save computational resources and time by focusing on most viable candidates. The phytocompounds were subjected to virtual screening to determine their drug-likeness and ADMET (Absorption, Distribution, Metabolism, Excretion, and Toxicity) properties, in accordance with the Lipinski’s rule of five. According to the rule of five, compounds are considered likely to be well absorbed when they possess partition coefficient (LogP) value less than 5, molecular weight less than 500, the number of hydrogen bond donors less than 5, the number of hydrogen bond acceptors less than 10, and the number of rotatable bonds less than 10 (Lipinski *et al*., 1997; Lipinski, 2004). Other parameters screened for were mutagenicity, carcinogenicity, reproductive effectiveness, ligand efficiency, drug-likeness, and irritancy (Table 3). These parameters allow screening out compounds that do not meet the physicochemical criteria for drug-like behavior (supplementary data). The 2D structures of statins were also downloaded from PubChem database and subjected to the same screening to serve as reference (Table 4).

**Table 3:** Drug-likeness and ADMET properties of top-ranked phytochemicals of *C. planchonii* and *C. tinctorium* at HMG-binding site of HMGR.

| S/N | Pubchem ID | Compound | MW (g/mol) | HA | HD | C logP | TPSA | DL | LE | RE | Mutagenic | Tumorigenic | Irritant |
| --- | --- | --- | --- | --- | --- | --- | --- | --- | --- | --- | --- | --- | --- |
| 1 | 73641 | Arjunolic acid | 488.71 | 5 | 4 | 4.286 | 97.99 | -5.555 | 0.162 | None | None | None | None |
| 2 | 12305768 | Alphitolic acid | 472.71 | 4 | 3 | 5.519 | 77.76 | -21.780 | 0.077 | None | None | None | None |
| 3 | 5281855 | Ellagic acid | 302.19 | 8 | 4 | 1.277 | 133.52 | -1.598 | 0.142 | None | None | None | None |
| 4 | 72277 | Epigallocatechin | 306.26 | 7 | 6 | 1.163 | 130.61 | 0.315 | 0.258 | None | None | None | None |
| 5 | 13915428 | 3-O-methylellagic acid | 316.22 | 8 | 3 | 1.553 | 122.52 | -1.390 | 0.111 | None | None | None | None |
| 6 | 5280417 | 3,7-di-O-methylquercetin | 330.29 | 7 | 3 | 2.194 | 105.45 | -0.105 | 0.130 | None | None | None | None |
| 7 | 44446550 | 3,4'-O-dimethylquercetin | 332.31 | 7 | 3 | 1.662 | 105.45 | 0.503 | 0.077 | None | None | None | None |
| 8 | 9064 | Catechin | 290.27 | 6 | 5 | 1.509 | 110.38 | 0.315 | 0.329 | None | None | None | None |
| 9 | 535203 | 3-(Azepan-1-yl)-1,2-benzothiazole 1,1-dioxide | 264.35 | 4 | 0 | 2.478 | 58.12 | -1.176 | 0.249 | None | None | None | None |
| 10 | 101202074 | (2E,4E,6E,8E,10E,12E)-13-[(1S)-1-hydroxy-2,6,6-trimethyl-4-oxocyclohex-2-en-1-yl]-2,6,11-trimethyltrideca-2,4,6,8,10,12-hexaenoic acid | 396.53 | 4 | 2 | 6.038 | 74.60 | 0.101 | 0.047 | None | None | None | None |

\*MW: Molecular Weight; HA: Hydrogen Acceptor; HD: Hydrogen Donor; ClogP: Octanol-water Partition Coefficient; TPSA: Topological Polar Surface Area; DL: Drug-likeness; LE: Ligand Efficiency; RE: Reproductive Effectiveness.

**Table 4:** Drug-likeness and ADMET properties of statins at HMG-binding site of HMGR.

| S/N | Pubchem ID | Compound | MW (g/mol) | HA | HD | C log P | TPSA | DL | LE | RE | Mutagenic | Tumorigenic | Irritant |
| --- | --- | --- | --- | --- | --- | --- | --- | --- | --- | --- | --- | --- | --- |
| 1 | PDB ID: 117/obj01 | Atorvastatin (Co-crystallized Control) | 558.65 | 7 | 4 | 5.622 | 111.79 | 4.451 | - | High | None | None | None |
| 2 | 60823 | Atorvastatin | 558.65 | 7 | 4 | 5.622 | 111.79 | 4.451 | 0.141 | High | None | None | None |
| 3 | 64715 | Mevastatin (Compactin) | 390.52 | 5 | 1 | 3.626 | 72.83 | 0.578 | 0.205 | None | None | None | None |
| 4 | 446155 | Fluvastatin | 411.47 | 5 | 3 | 3.978 | 82.69 | 1.804 | 0.153 | High | None | None | None |
| 5 | 446156 | Cerivastatin | 459.56 | 6 | 3 | 4.320 | 99.88 | -0.283 | 0.139 | None | None | None | None |
| 6 | 446157 | Rosuvastatin | 481.54 | 9 | 3 | 2.100 | 149.3 | 3.454 | 0.139 | None | None | None | None |
| 7 | 54454 | Simvastatin | 418.57 | 5 | 1 | 4.461 | 72.83 | 0.668 | 0.195 | None | None | None | None |

\*MW: Molecular Weight; HA: Hydrogen Acceptor; HD: Hydrogen Donor; C log P: Octanol-water Partition Coefficient; TPSA: Topological Polar Surface Area; DL: Drug-likeness; LE: Ligand Efficiency; RE: Reproductive Effectiveness.

### Drug Target Preparation

The 3D crystal structure of human HMGR (PDB ID: 1HWK) complexed with atorvastatin (PDB ID: 117) was retrieved in PDB format 5^th^ August, 2024 from Protein Data Bank (PDB) (https://www.rcsb.org). The drug target was prepared by removing non-essential subunits (B, C, D), Adenosine diphosphate (ADP), heteroatoms, and water molecules using PyMOL visualization tool (https://www.pymol.org). The unique ligand atorvastatin (obj01/117), which served as one of the control ligands was extracted from the catalytic subunit A, in addition to the 6 other statins downloaded from PubChem. Both target and ligand were saved in PDB and SDF formats respectively. Using PyMOL allows one to visualize and predict the grid co-ordinates around the HMG-binding pocket, while Discovery Studio visualizer (https://www.3ds.com) helps in identifying and characterizing the residues at the binding site.

### Molecular Docking Analysis

PyRx virtual screening tool (https://pyrx.sourceforge.io) was used for the molecular docking. The prepared drug target HMGR was loaded on PyRx in PDB format, hydrogen atoms were added to ensure the protein is correctly protonated and made as macromolecule, after which the screened phytochemicals were imported in SDF format. These compounds were subjected to energy minimization using the optimization algorithm tool of PyRx and the required force field was set at “ghemical”, adjusting the positions of atoms in the phytochemicals in order to reduce their overall energy and optimize them. After their conversion to PDBQT format, the 3D grid box which encloses the HMG-binding pocket (residues 682**-**694) of the protein, where the compounds will bind was centered at co-ordinates 22.2175 × -3.5559 × 5.8150 along X, Y, and Z axes respectively, at grid dimensions 21.0454 × 28.2041 × 28.7731 Å along the same axes. This type of docking is semi-rigid, where the structure of receptor (HMGR) remains rigid while the phytochemicals and statins have some degree of flexibility at the binding pocket. In the analysis, the PyRx AutoDock Vina Wizard exhaustive search docking function was used for the molecular docking. To ensure the feasibility and validity of the docking study, the native ligand atorvastatin (obj01/117 extracted from the drug target 1HWK) and 6 other statins including atorvastatin itself, were downloaded from PubChem. Their interactions in the respective HMGR complexes (PDB IDs: 1HWK, 1HWI, 1HWL, 1HWJ, 1HW8, 1HW9, https://www.rcsb.org), as reported by Istvan and Deisenhofer, were initially examined. The statins were then docked at the HMG-binding site, to ascertain the accuracy of the PyRx algorithm. Their results were compared with those in the literature, after which the docking of the 32 hit compounds was performed. The results were exported as PDBQT files to evaluate the best poses (binding modes) of the protein-ligand complexes using PyMOL and Discovery Studio tools. Their binding energy scores were saved in excel format for further analysis. The docking process was repeated for all 84 phytochemicals, without screening for their drug-likeness and toxicity properties, to determine whether potential inhibitors, which might have been previously screened out, could be identified as drug candidates.

## Results

In order to investigate the mechanism of binding and inhibition of bioactive compounds isolated from *C. planchonii* and *C. tinctorium* on human HMGR activity as potential cholesterol-lowering agents, statins and each compound were docked against the HMG-binding pocket (residues 682**-**694) of the enzyme. The docking study results revealed that 10 lead compounds exhibited strong binding affinities, with binding energy (ΔG) scores ranging from -4.6 to -6.0 kcal/mol (Fig. 3; Table 5). These phytochemicals also interacted well with the relevant amino acid residues at the HMG-binding pocket of the enzyme (Fig. 4). Their ΔG scores were comparable to or exceeded those of the control ligands (-4.6 to -5.7 kcal/mol; Table 6). Their docking scores also compared favorably with those of statins. One of the lead compounds, 3-O-methylellagic acid (ID_13915428) demonstrated stronger and more substantial binding interactions with the HMG-binding pocket of the drug target than any other compounds, including statins, in addition to exhibiting high binding energy (Table 8a).

**Figure 3:**
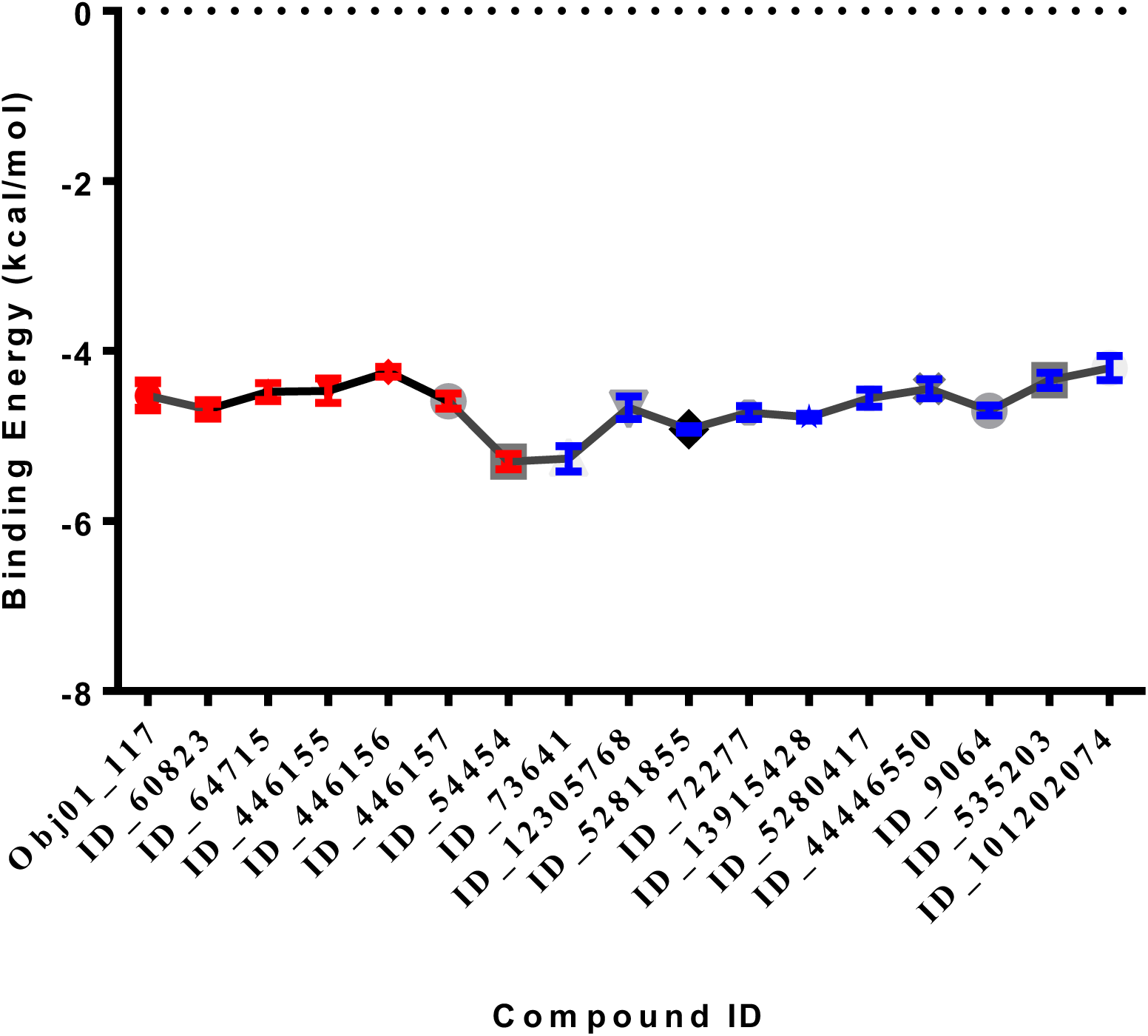
Binding potential of statins (red) and top-ranked phytochemicals (blue) at HMG-binding pocket of human HMGR (PDB ID: 1HWK). Values expressed are means ± SEM of 9 replicates at different binding poses. All phytochemicals represented satisfy Lipinski’s rule of 5.

**Table 5:**
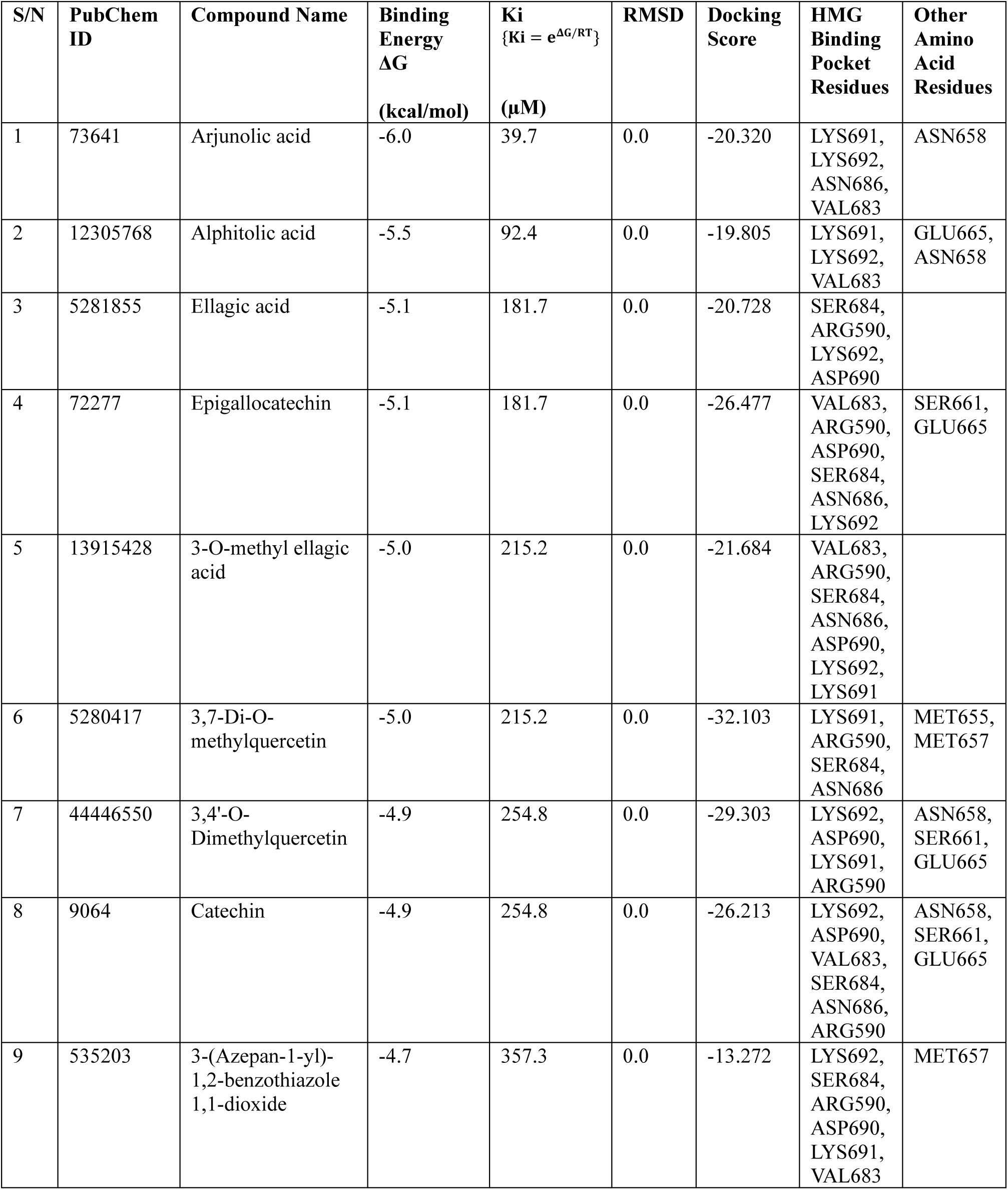
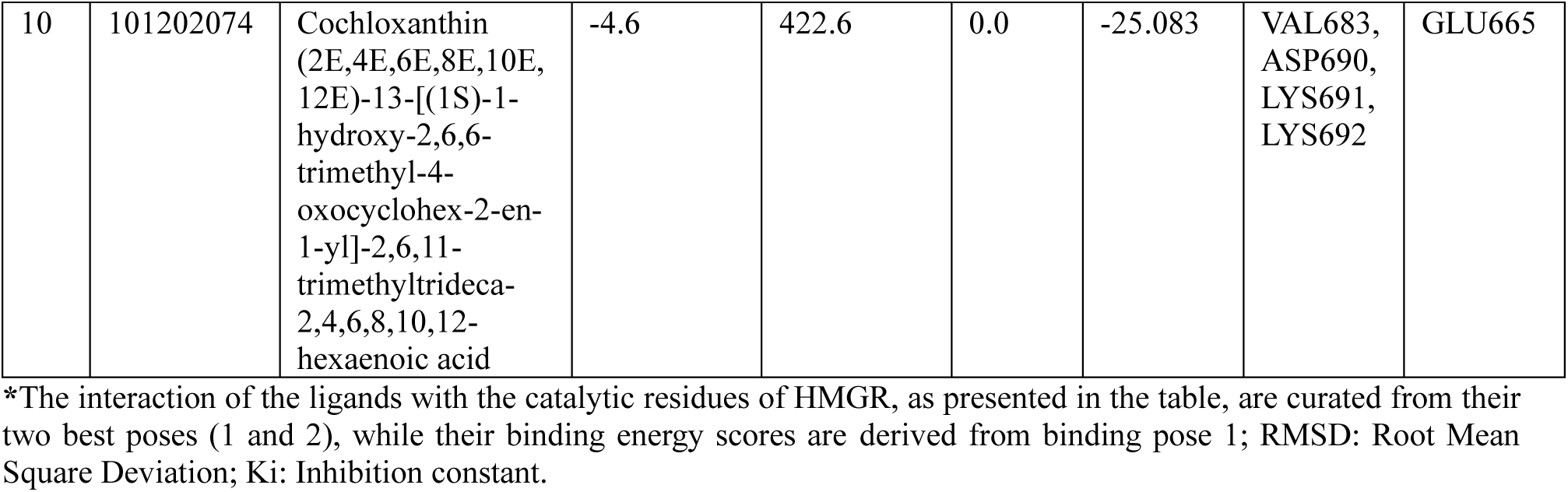
Molecular docking results of the top-ranked phytochemicals-binding at HMG-binding pocket of HMGR (PDB ID: 1HWK)

**Figure 4:**
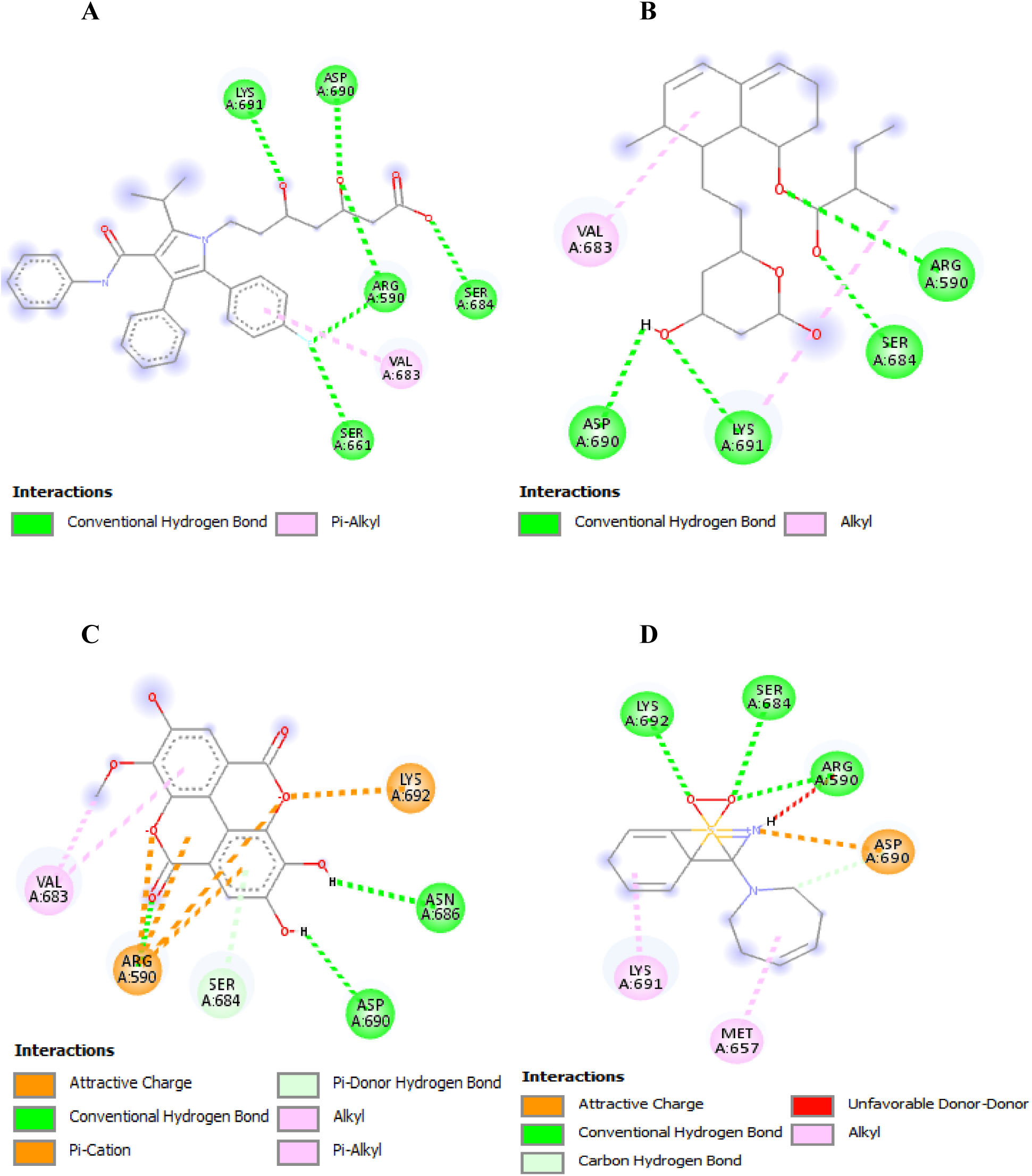
2D representation showing the binding interaction of **(A)** native atorvastatin (PDB ID_117) from literature, **(B)** mevastatin (ID_64715) pose 1, **(C**) ID_13915428 pose 1 and **(D)** ID_535203 pose 1, with HMG-binding pocket residues of human HMGR.

**Table 6.**
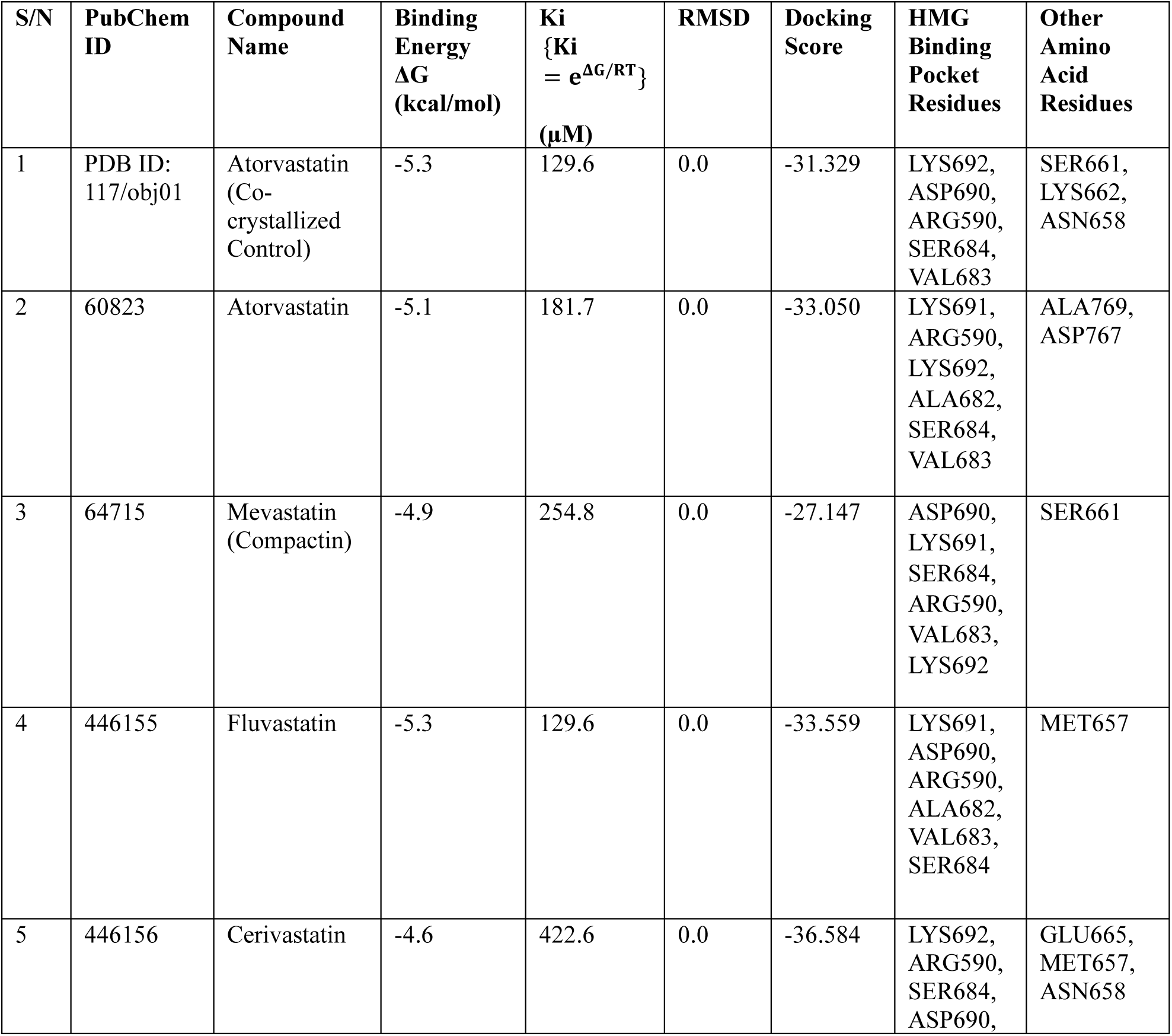
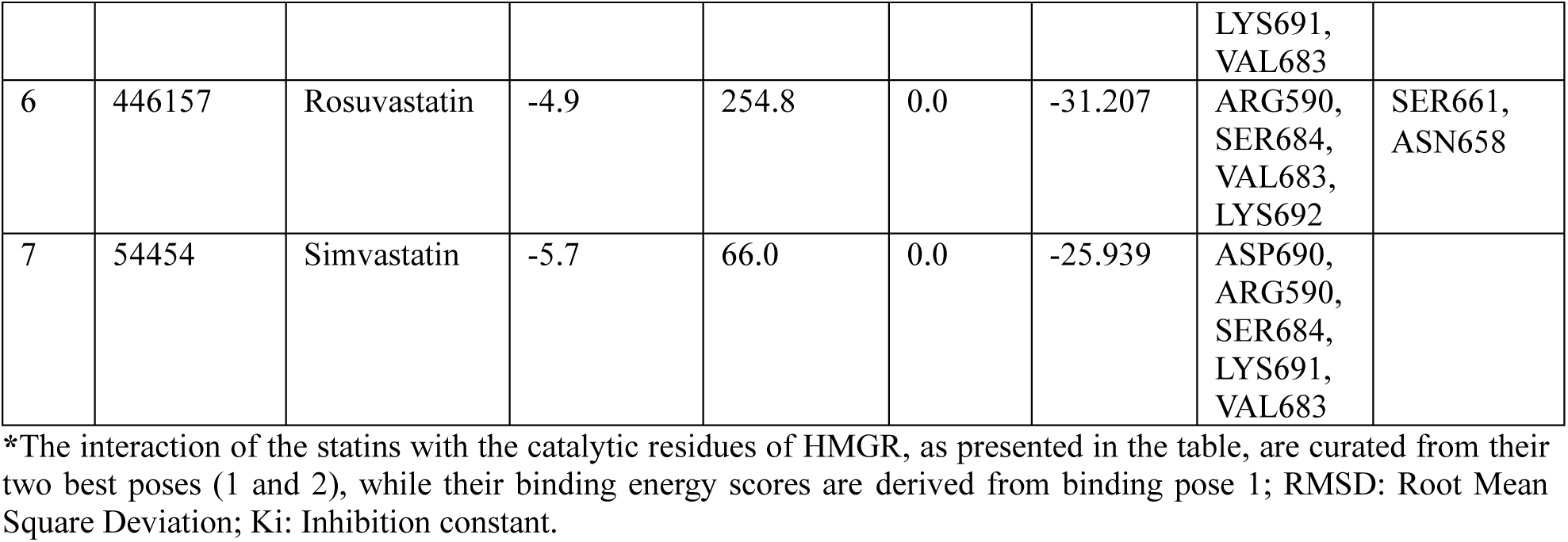
Molecular docking results of statins-binding at HMG-binding pocket of HMGR (PDB ID: 1HWK)

The docking score and binding energy score are two key metrics used in this molecular docking study (Table 5; 6). The docking score was a value generated by DataWarrior software to represent the quality of the ligand’s fit into the receptor’s binding site, and is derived using scoring function based on factors such as hydrogen bonding, van der Waal’s, hydrophobic, and electrostatic interactions (Sander *et al*., 2015). The docking score was mainly used to rank the different compounds in terms of how well they bind to the active site of HMGR and to compare the quality and fitness of their binding with those of statins within the same docking study. Higher docking scores (more negative) generally indicate a better fit between the ligand and drug target and vice versa. However, the docking score is not an absolute energy value.

In contrast, the binding energy score, represented as the free energy of binding (ΔG), measured in kcal/mol, refers to the affinity and strength of the interaction between a ligand and its target. It was generated by the AutoDock Vina algorithm in PyRx. The binding energy score predicts how strongly a ligand will bind to the target under physiological conditions, with higher (more negative) values indicating a stronger binding and more thermodynamically favorable formation of complexes (Trott and Olson 2010). Unlike docking score, the binding energy score was directly correlated with inhibition constant (Ki) values using the formula {Ki = e^ΔG/RT^}, where R is molar gas constant (1.987 cal/mol/K), and T is standard temperature in Kelvin (298K). Therefore, the selection of compounds was focused on those with stronger interactions and more effective binding energies rather than the ones with good docking scores, in addition to their drug-likeness properties.

As shown in Table 3 and 4, the drug-likeness score is also a crucial parameter used in determining whether a compound is likely to be an effective drug. A positive score indicates that a compound possesses structural features similar to known drugs, while a negative score suggests that such compound has structural features that are less common in known drugs. Good drug-like compounds usually have scores greater than zero (Sander *et al*., 2015; Lipinski, 2004). Ligand efficiency (LE) is a metric used to evaluate the binding efficiency of compounds relative to their size. A higher LE score indicates that a compound achieves its binding affinity with fewer atoms making it more efficient, while a lower LE score suggests that a compound relies on a larger structure to achieve its binding, which might be less desirable (Hopkins *et al*., 2014). When screening for toxic compounds, those that may bind to unintended off-target sites which could lead to adverse effects such as genetic mutations contributing to cancer development, or cause irritation to tissues like skin, eyes, or mucous membrane, were eliminated. Reproductive effectiveness parameter was used to predict the potential impact of a compound on reproductive health, including infertility and harm to fetal development (Sander *et al*., 2015).

## Discussion of Results

As shown in table 5, the 10 top-ranked phytochemicals identified in this study, in no particular order, comprise 2 hydrolysable tannins (ellagitannins): ellagic acid (ID_5281855) and 3-O-methylellagic acid (ID_13915428); 4 flavonoids: catechin (ID_9064), epigallocatechin (ID_72277), 3,7-Di-O-methylquercetin (ID_5280417), and 3,4’-O-Dimethylquercetin (ID_44446550); 2 triterpenoid saponins: arjunolic acid (ID_73641) and alphitolic acid (ID_12305768); 1 carotenoid: cochloxanthin (ID_101202074); and a benzothiazole derivative, 3-(Azepan-1-yl)-1,2-benzothiazole 1,1-dioxide (ID_535203). The molecular docking analyses of the two best binding modes of these compounds demonstrated their cholesterol-lowering potential, as they clearly show strong biochemical interactions and high binding energies with the relevant amino acid residues that constitute the HMG-binding pocket (cis-loop) of HMGR (Table 8), similar to or better than statins (Table 7). This suggests that they could hinder the binding of the substrate HMG-CoA through competitive inhibition, similar to statins.

**Table 7a.** Interaction profile of 1172 at HMG-binding site of HMGR (Istvan and Deisenhofer, 2001).

| HMGR-Statin Interaction | Bond Distance | Category |
| --- | --- | --- |
| A:ARG590:NE - A:1172:F1 | 2.76608 | Hydrogen Bond; Halogen |
| A:ARG590:NH1 - A:1172:O3 | 3.02765 | Hydrogen Bond |
| A:ARG590:NH2 - A:1172:O3 | 3.27471 | Hydrogen Bond |
| A:SER661:OG - A:1172:F1 | 3.27271 | Hydrogen Bond; Halogen |
| A:SER684:OG - A:1172:O1A | 2.65383 | Hydrogen Bond |
| A:LYS691:NZ - A:1172:O5 | 2.84259 | Hydrogen Bond |
| A:1172:O3 - A:ASP690:OD2 | 2.79251 | Hydrogen Bond |
| A:1172 - A:VAL683 | 5.18646 | Hydrophobic |

\*1172: Native/co-crystallized atorvastatin; A: Amino Acid.

**Table 7b.** Interaction profile of ID_117/obj01 at HMG-binding site of HMGR.

| HMGR-Statin Interaction | Bond Distance | Category |
| --- | --- | --- |
| A:SER661:HG - N:UNK1:O | 2.43435 | Hydrogen Bond |
| A:LYS692:HZ2 - N:UNK1:O | 2.2243 | Hydrogen Bond |
| N:UNK1:H - A:ASP690:O | 2.75222 | Hydrogen Bond |
| N:UNK1:H - A:ASP690:OD2 | 2.08098 | Hydrogen Bond |
| N:UNK1:H - N:UNK1:O | 2.31052 | Hydrogen Bond |
| A:ASN658:CA - N:UNK1:O | 3.66311 | Hydrogen Bond |
| A:SER684:CB - N:UNK1:O | 3.5454 | Hydrogen Bond |
| A:ARG590:NH2 - N:UNK1 | 4.24896 | Electrostatic |
| N:UNK1:C - A:VAL683 | 4.43649 | Hydrophobic |
| N:UNK1 - A:LYS662 | 5.34202 | Hydrophobic |
| A:LYS692:HZ2 - N:UNK1:O | 2.42944 | Hydrogen Bond |
| N:UNK1:H - A:SER684:O | 2.3963 | Hydrogen Bond |
| N:UNK1:H - A:ASP690:OD2 | 2.77571 | Hydrogen Bond |
| A:ARG590:NH2 - N:UNK1 | 3.99181 | Electrostatic |

\*UNK1: Native/co-crystallized atorvastatin; A: Amino Acid.

**Table 7c.** Interaction profile of ID_60823 at HMG-binding site of HMGR.

| HMGR-Statin Interaction | Bond Distance | Category |
| --- | --- | --- |
| A:ARG590:HH11 - N:UNK1:O | 2.32911 | Hydrogen Bond |
| A:ARG590:HH21 - N:UNK1:O | 2.30312 | Hydrogen Bond |
| A:LYS691:HZ3 - N:UNK1:O | 2.19002 | Hydrogen Bond |
| A:LYS692:HZ1 - N:UNK1:O | 2.70078 | Hydrogen Bond |
| N:UNK1:H - N:UNK1:O | 1.90918 | Hydrogen Bond |
| A:ALA769:CB - N:UNK1 | 3.94942 | Hydrophobic |
| N:UNK1 - N:UNK1 | 5.10611 | Hydrophobic |
| N:UNK1 - A:LYS691 | 4.66065 | Hydrophobic |
| N:UNK1 - A:ALA769 | 4.52421 | Hydrophobic |
| N:UNK1 - A:LYS692 | 4.22164 | Hydrophobic |
| N:UNK1 - A:LYS691 | 4.96112 | Hydrophobic |
| A:ARG590:HH11 - N:UNK1:O | 2.48921 | Hydrogen Bond |
| A:ARG590:HH22 - N:UNK1:O | 2.50246 | Hydrogen Bond |
| A:SER661:HG - N:UNK1:O | 2.69997 | Hydrogen Bond |
| A:SER684:HG - N:UNK1:O | 2.00339 | Hydrogen Bond |
| A:LYS692:HZ1 - N:UNK1:O | 2.74314 | Hydrogen Bond |
| A:LYS692:HZ2 - N:UNK1:O | 2.35487 | Hydrogen Bond |
| A:LYS692:HZ3 - N:UNK1:O | 2.98317 | Hydrogen Bond |
| N:UNK1:H - A:ASP690:O | 2.33486 | Hydrogen Bond |
| N:UNK1:H - A:ASP690:OD2 | 3.03387 | Hydrogen Bond |
| N:UNK1:H - A:ASN658:OD1 | 1.68671 | Hydrogen Bond |
| A:ARG590:NH2 - N:UNK1 | 4.01054 | Electrostatic |
| A:VAL683:CG1 - N:UNK1 | 3.93163 | Hydrophobic |

\*UNK1: Atorvastatin; A: Amino Acid.

**Table 7d.** Interaction profile of ID_54454 at HMG-binding site of HMGR.

| HMGR-Statin Interaction | Bond Distance | Category |
| --- | --- | --- |
| A:ARG590:HH11 - N:UNK1:O | 2.16123 | Hydrogen Bond |
| A:ARG590:HH21 - N:UNK1:O | 2.72168 | Hydrogen Bond |
| A:ARG590:HH21 - N:UNK1:O | 2.12832 | Hydrogen Bond |
| A:ARG590:HH21 - N:UNK1:O | 2.7818 | Hydrogen Bond |
| A:SER684:HG - N:UNK1:O | 2.18719 | Hydrogen Bond |
| N:UNK1:H - A:ASP690:OD2 | 2.02284 | Hydrogen Bond |
| N:UNK1:C - A:LYS691 | 3.7506 | Hydrophobic |
| A:ARG590:HH11 - N:UNK1:O | 2.07764 | Hydrogen Bond |
| A:ARG590:HH21 - N:UNK1:O | 2.42845 | Hydrogen Bond |
| A:SER684:HG - N:UNK1:O | 2.70634 | Hydrogen Bond |
| N:UNK1:H - A:ASP690:OD2 | 2.57367 | Hydrogen Bond |
| N:UNK1:H - N:UNK1:O | 1.85958 | Hydrogen Bond |
| A:VAL683 - N:UNK1 | 5.13668 | Hydrophobic |
| N:UNK1:C - A:LYS691 | 3.70484 | Hydrophobic |

\*UNK1: Simvastatin; A: Amino Acid.

**Table 7e.** Interaction profile of ID_446155 at HMG-binding site of HMGR.

| HMGR-Statin Interaction | Bond Distance | Category |
| --- | --- | --- |
| A:ARG590:HH21 - N:UNK1:O | 2.04568 | Hydrogen Bond |
| A:LYS691:HZ1 - N:UNK1:O | 2.76556 | Hydrogen Bond |
| A:LYS691:HZ3 - N:UNK1:O | 1.92611 | Hydrogen Bond |
| A:LYS691:HZ3 - N:UNK1:O | 2.96588 | Hydrogen Bond |
| N:UNK1:H - A:ASP690:OD2 | 2.44339 | Hydrogen Bond |
| A:VAL683:CG1 - N:UNK1 | 3.93996 | Hydrophobic |
| A:ALA682 - N:UNK1 | 4.68522 | Hydrophobic |
| A:LYS692:HZ2 - N:UNK1:F | 2.70465 | Hydrogen Bond; Halogen |
| A:LYS692:HZ3 - N:UNK1:F | 2.45578 | Hydrogen Bond; Halogen |
| N:UNK1:H - A:SER684:O | 2.59951 | Hydrogen Bond |
| A:SER684:CB - N:UNK1:O | 3.36541 | Hydrogen Bond |
| A:MET657 - N:UNK1 | 4.69986 | Hydrophobic |

\*UNK1: Fluvastatin; A: Amino Acid.

**Table 7f.** Interaction profile of ID_446157 at HMG-binding site of HMGR.

| HMGR-Statin Interaction | Bond Distance | Category |
| --- | --- | --- |
| A:ARG590:HH22 - N:UNK1:O | 2.10885 | Hydrogen Bond |
| A:SER661:HG - N:UNK1:O | 2.01271 | Hydrogen Bond |
| A:SER684:HG - N:UNK1:O | 2.62883 | Hydrogen Bond |
| N:UNK1:H - A:ASN658:OD1 | 2.05769 | Hydrogen Bond |
| N:UNK1:HN - N:UNK1:O | 2.46718 | Hydrogen Bond |
| N:UNK1:C - N:UNK1:F | 3.60226 | Halogen |
| N:UNK1 - A:VAL683 | 4.39623 | Hydrophobic |
| A:ARG590:HH22 - N:UNK1:O | 2.32736 | Hydrogen Bond |
| A:SER661:HG - N:UNK1:O | 2.43806 | Hydrogen Bond |
| A:SER661:HG - N:UNK1:O | 2.13 | Hydrogen Bond |
| A:LYS692:HZ2 - N:UNK1:N | 2.83329 | Hydrogen Bond |
| A:LYS692:HZ3 - N:UNK1:O | 1.98837 | Hydrogen Bond |
| N:UNK1:HN - N:UNK1:O | 2.48714 | Hydrogen Bond |
| A:ASN658:CA - N:UNK1:O | 3.68275 | Hydrogen Bond |
| N:UNK1:C - N:UNK1 | 3.88326 | Hydrophobic |
| N:UNK1:C - N:UNK1 | 3.89806 | Hydrophobic |

\*UNK1: Rosuvastatin; A: Amino Acid.

**Table 7g.**
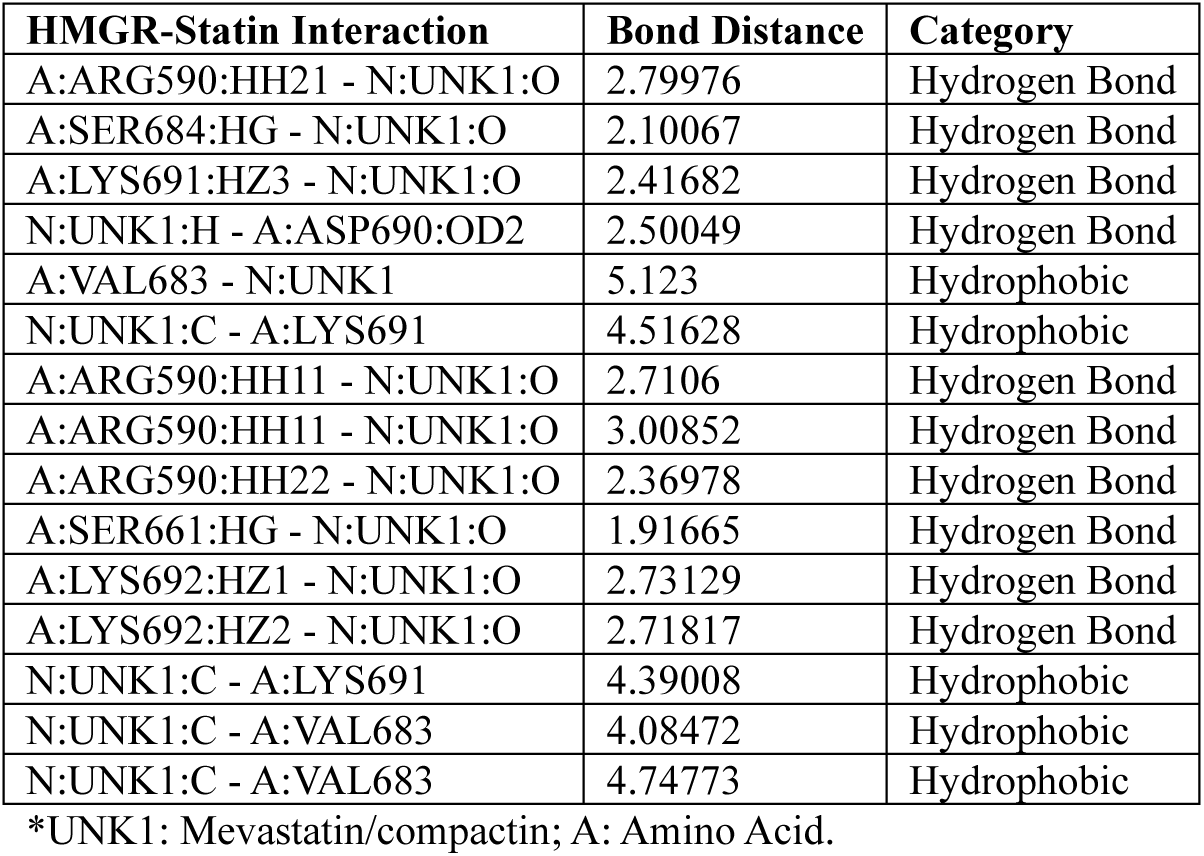
Interaction profile of ID_64715 at HMG-binding site of HMGR.

**Table 7h.** Interaction profile of ID_446156 at HMG-binding site of HMGR.

| HMGR-Statin Interaction | Bond Distance | Category |
| --- | --- | --- |
| A:SER684:HG - N:UNK1:O | 2.06156 | Hydrogen Bond |
| N:UNK1:H - A:ASP690:O | 2.37362 | Hydrogen Bond |
| N:UNK1:C - A:GLU665:OE2 | 3.65532 | Hydrogen Bond |
| N:UNK1:C - A:MET657 | 4.50015 | Hydrophobic |
| N:UNK1:N - A:GLU665:OE2 | 4.3182 | Electrostatic |
| A:ARG590:HH11 - N:UNK1:O | 2.52497 | Hydrogen Bond |
| A:ARG590:HH21 - N:UNK1:O | 2.09354 | Hydrogen Bond |
| A:LYS691:HZ3 - N:UNK1:O | 2.83346 | Hydrogen Bond |
| N:UNK1:H - A:ASN658:OD1 | 2.27593 | Hydrogen Bond |
| N:UNK1:C - A:GLU665:OE2 | 3.67977 | Hydrogen Bond |
| N:UNK1:C - N:UNK1 | 3.77546 | Hydrophobic |
| N:UNK1:C - A:VAL683 | 4.29239 | Hydrophobic |
| N:UNK1:C - A:VAL683 | 4.28785 | Hydrophobic |

\*UNK1: Cerivastatin; A: Amino Acid.

Several polar interactions with the cis-loop residues (Arg^590^, Ser^684^, Asn^686^, Asp^690^, Lys^691^, Lys^692^) of HMGR, are formed by the hydroxyl (-OH) groups of the aromatic rings, carbonyl groups (C=O), and lactone ring oxygen atoms of the ellagitannins (ID_5281855 and ID_13915428). Their bulky rings also establish several electrostatic and hydrophobic contacts with residues Val^683^, Arg^590^, Ser^684^, Asp^690^, Lys^691^, and Lys^692^ (Table 8a-b; Fig. 3C; supplementary data). No interactions of these polyphenols were observed with other residues within the HMGR binding site. Among all the compounds, including statins, 3-O-methylellagic acid (ID_13915428) exhibits the greatest number (26) of binding interactions (Table 8a), indicating that this polyphenolic compound could be a viable drug candidate for HMGR inhibition. A recent in vivo and in vitro study by Lee et al. demonstrated that ellagic acid inhibits HMGR by activating AMP-activated protein kinase (AMPK), leading to the phosphorylation and inactivation of the enzyme. This study, which included rats fed a high-cholesterol diet, revealed that the administration of ellagic acid (4 mg/kg/day, p.o) resulted in significant reductions in serum total cholesterol, LDL-C, and triglyceride levels. It was also found to downregulate the gene expression of sterol regulatory element-binding protein-2 (SREBP-2) and its target protein HMG-CoA reductase (HMGR), thereby reducing cholesterol biosynthesis in the liver (Lee *et al*., 2020). In addition to its roles in cholesterol metabolism, ellagic acid and its derivatives exhibit antioxidant and anti-inflammatory properties, which contribute to their protective effects against cardiovascular diseases (Salvamani *et al*., 2016).

**Table 8a.** Interaction profile of ID_13915428 at HMG-binding site of HMGR.

| HMGR-Phytochemical Interaction | Bond Distance | Category |
| --- | --- | --- |
| A:ARG590:NH1 - N:UNK1:O | 5.44885 | Electrostatic |
| A:ARG590:NH2 - N:UNK1:O | 3.84437 | Electrostatic |
| A:LYS692:NZ - N:UNK1:O | 5.55951 | Electrostatic |
| A:ARG590:HH21 - N:UNK1:O | 2.38275 | Hydrogen Bond |
| N:UNK1:H - A:ASN686:OD1 | 2.65517 | Hydrogen Bond |
| N:UNK1:H - A:ASP690:O | 2.43114 | Hydrogen Bond |
| A:ARG590:NH1 - N:UNK1 | 4.31545 | Electrostatic |
| A:ARG590:NH1 - N:UNK1 | 3.91213 | Electrostatic |
| A:SER684:HG - N:UNK1 | 3.1568 | Hydrogen Bond |
| N:UNK1:C - A:VAL683 | 4.87718 | Hydrophobic |
| N:UNK1 - A:VAL683 | 5.3518 | Hydrophobic |
| A:ARG590:NH1 - N:UNK1:O | 3.52478 | Electrostatic |
| A:ARG590:NH2 - N:UNK1:O | 4.71756 | Electrostatic |
| A:LYS691:NZ - N:UNK1:O | 4.63929 | Electrostatic |
| A:SER684:HG - N:UNK1:O | 2.10052 | Hydrogen Bond |
| A:LYS691:HZ3 - N:UNK1:O | 1.90064 | Hydrogen Bond |
| A:LYS692:HZ3 - N:UNK1:O | 2.54599 | Hydrogen Bond |
| N:UNK1:H - A:ASP690:O | 2.16329 | Hydrogen Bond |
| A:SER684:CB - N:UNK1:O | 3.56971 | Hydrogen Bond |
| A:SER684:CB - N:UNK1:O | 3.52304 | Hydrogen Bond |
| N:UNK1:C - A:ASN686:OD1 | 3.55315 | Hydrogen Bond |
| A:ARG590:NH1 - N:UNK1 | 4.12152 | Hydrogen Bond; Electrostatic |
| A:ARG590:NH2 - N:UNK1 | 3.53239 | Electrostatic |
| A:ARG590:NH2 - N:UNK1 | 3.87267 | Electrostatic |
| A:ARG590:NH2 - N:UNK1 | 4.12093 | Hydrogen Bond; Electrostatic |
| A:ASP690:OD2 - N:UNK1 | 3.86666 | Electrostatic |

\*UNK1: 3-O-methylelagic acid; A: Amino acid.

**Table 8b.** Interaction profile of ID_5281855 at HMG-binding site of HMGR.

| HMGR-Phytochemical Interaction | Bond Distance | Category |
| --- | --- | --- |
| A:ARG590:HH21 - N:UNK1:O | 2.81417 | Hydrogen Bond; Electrostatic |
| A:LYS692:NZ - N:UNK1:O | 4.98035 | Electrostatic |
| A:ARG590:HH21 - N:UNK1:O | 2.61024 | Hydrogen Bond |
| A:LYS692:HZ3 - N:UNK1:O | 2.67545 | Hydrogen Bond |
| N:UNK1:H - A:ASP690:O | 2.69978 | Hydrogen Bond |
| N:UNK1:H - A:ASP690:OD2 | 2.13329 | Hydrogen Bond |
| A:ARG590:NH1 - N:UNK1 | 4.157 | Electrostatic |
| A:ARG590:NH1 - N:UNK1 | 3.84096 | Electrostatic |
| A:SER684:CB - N:UNK1 | 3.27967 | Hydrophobic |
| A:ARG590:HH21 - N:UNK1:O | 2.87531 | Hydrogen Bond; Electrostatic |
| A:LYS692:NZ - N:UNK1:O | 4.95449 | Electrostatic |
| A:LYS692:HZ3 - N:UNK1:O | 2.64235 | Hydrogen Bond |
| N:UNK1:H - A:ASP690:OD2 | 2.11316 | Hydrogen Bond |
| N:UNK1:H - A:SER684:O | 2.57614 | Hydrogen Bond |
| A:ARG590:NH1 - N:UNK1 | 3.87895 | Electrostatic |
| A:ARG590:NH1 - N:UNK1 | 4.17645 | Electrostatic |
| A:SER684:CB - N:UNK1 | 3.30588 | Hydrophobic |

\*UNK1: Ellagic acid; A: Amino acid.

The four flavonoids identified, belonging to the catechin and quercetin classes of polyphenols (ID_9064, ID_72277, ID_5280417, ID_44446550), demonstrated their potential to mimic the binding of statins by forming polar hydrogen bonds with cis-loop residues (Arg^590^, Ser^684^, Asn^686^, Asp^690^, Lys^691^, Lys^692^) and other residues (Asn^658^, Glu^665^). They also formed several electrostatic and non-polar hydrophobic interactions with Val^683^ and other residues, including Met^655^, Met^657^, and Ser^661^, at the HMGR active site. This capability is attributed to their basic flavan-ring structure with multiple polar -OH, C=O, pyran ring oxygen atoms, and methoxy (-OCH3) groups (Table 8c-f; supplementary data). An in vitro experiment showed that catechin isolates from *Uncaria gambir*, an Indonesian plant, exhibit strong inhibitory activities against HMGR with 97.46% efficacy, compared to 85.74% for simvastatin, a performance suggesting it could stand out as a promising therapy for hypercholesterolemia treatment (Yunarto *et al*., 2021). Surprisingly, epigallocatechin gallate (EGCG) has been shown to potently and reversibly inhibit HMGR in vitro by competing with its cofactor NADPH and binding at the cofactor site instead of the HMG-binding pocket (Cuccioloni *et al*., 2011). However, this present study suggests that EGCG may possess both capabilities. Quercetin dihydrate and gallate supplements have also been reported to lower plasma and hepatic cholesterol levels in rats fed a cholesterol-rich diet. The results of the study concluded that quercetin dihydrate significantly reduced hepatic HMGR activity compared to normal control groups (Bok *et al*., 2002). Furthermore, several other studies have elucidated on the ability of quercetin to drastically reduce HMGR activity, inhibit fatty acid and triacylglycerol synthesis in hepatocytes, and alleviate endothelial dysfunction associated with age-related cardiovascular diseases (Khamis *et al*., 2017; Gnoni *et al*., 2009; Dagher *et al*., 2021).

**Table 8c.** Interaction profile of ID_9064 at HMG-binding site of HMGR.

| HMGR-Phytochemical Interaction | Bond Distance | Category |
| --- | --- | --- |
| A:LYS692:HZ2 - N:UNK1:O | 2.21812 | Hydrogen Bond |
| N:UNK1:H - A:ASP690:O | 2.69834 | Hydrogen Bond |
| N:UNK1:H - A:ASN686:OD1 | 2.23274 | Hydrogen Bond |
| N:UNK1:H - A:ASN658:OD1 | 2.51386 | Hydrogen Bond |
| A:SER661:HG - N:UNK1 | 3.22519 | Hydrogen Bond |
| A:SER684:HG - N:UNK1:O | 2.26898 | Hydrogen Bond |
| N:UNK1:H - A:GLU665:OE2 | 2.15329 | Hydrogen Bond |
| A:SER661:CB - N:UNK1:O | 3.28108 | Hydrogen Bond |
| A:ARG590:NH1 - N:UNK1 | 3.79509 | Electrostatic |
| A:VAL683:CG1 - N:UNK1 | 3.71449 | Hydrophobic |

\*UNK1: Catechin; A: Amino acid.

**Table 8d.** Interaction profile of ID_72277 at HMG-binding site of HMGR.

| HMGR-Phytochemical Interaction | Bond Distance | Category |
| --- | --- | --- |
| A:LYS692:HZ1 - N:UNK1:O | 2.59154 | Hydrogen Bond |
| N:UNK1:H - A:ASP690:O | 2.08395 | Hydrogen Bond |
| N:UNK1:H - A:GLU665:OE2 | 2.01012 | Hydrogen Bond |
| A:SER661:CB - N:UNK1:O | 3.22728 | Hydrogen Bond |
| A:ARG590:NH1 - N:UNK1 | 3.745 | Electrostatic |
| A:VAL683:CG1 - N:UNK1 | 3.63107 | Hydrophobic |
| A:LYS692:HZ2 - N:UNK1:O | 2.86521 | Hydrogen Bond |
| N:UNK1:H - A:ASP690:O | 2.11276 | Hydrogen Bond |
| N:UNK1:H - A:ASN686:OD1 | 3.0048 | Hydrogen Bond |
| A:ARG590:NH1 - N:UNK1 | 3.71072 | Electrostatic |
| A:VAL683:CG1 - N:UNK1 | 3.72652 | Hydrophobic |

\*UNK1: Epigallocatechin; A: Amino acid.

**Table 8e.** Interaction profile of ID_5280417 at HMG-binding site of HMGR.

| HMGR-Phytochemical Interaction | Bond Distance | Category |
| --- | --- | --- |
| A:SER684:HG - N:UNK1:O | 1.9721 | Hydrogen Bond |
| A:LYS691:HZ1 - N:UNK1:O | 2.86506 | Hydrogen Bond |
| A:ARG590:NH2 - N:UNK1 | 3.37979 | Electrostatic |
| A:ARG590:NH2 - N:UNK1 | 4.20141 | Electrostatic |
| N:UNK1:C - A:MET655 | 5.33201 | Hydrophobic |
| N:UNK1:C - A:MET657 | 5.06009 | Hydrophobic |
| N:UNK1 - A:MET657 | 5.46588 | Hydrophobic |
| N:UNK1 - A:MET657 | 4.66849 | Hydrophobic |
| A:ARG590:HH21 - N:UNK1:O | 2.84211 | Hydrogen Bond |
| A:LYS691:HZ3 - N:UNK1:O | 2.17156 | Hydrogen Bond |
| A:LYS691:HZ3 - N:UNK1:O | 2.35692 | Hydrogen Bond |
| N:UNK1:H - A:ASN686:OD1 | 2.07754 | Hydrogen Bond |
| A:ARG590:NH2 - N:UNK1 | 4.1918 | Hydrogen Bond; Electrostatic |

\*UNK1: 3,7-Di-O-methylquercetin; A: Amino acid.

**Table 8f.** Interaction profile of ID_44446550 at HMG-binding site of HMGR.

| HMGR-Phytochemical Interaction | Bond Distance | Category |
| --- | --- | --- |
| A:LYS692:HZ2 - N:UNK1:O | 2.37781 | Hydrogen Bond |
| A:LYS692:HZ3 - N:UNK1:O | 2.24752 | Hydrogen Bond |
| N:UNK1:H - A:ASN658:OD1 | 2.54886 | Hydrogen Bond |
| N:UNK1:H - A:GLU665:OE2 | 2.13089 | Hydrogen Bond |
| N:UNK1:C - A:ASP690:OD2 | 3.68626 | Hydrogen Bond |
| A:ARG590:NH2 - N:UNK1 | 3.31539 | Electrostatic |
| A:ARG590:NH2 - N:UNK1 | 4.12013 | Electrostatic |
| A:SER661:HG - N:UNK1 | 3.05821 | Hydrogen Bond |
| A:SER661:HG - N:UNK1:O | 2.99324 | Hydrogen Bond |
| A:LYS691:HZ3 - N:UNK1:O | 2.04514 | Hydrogen Bond |
| A:LYS692:HZ1 - N:UNK1:O | 2.77091 | Hydrogen Bond |
| A:LYS692:HZ3 - N:UNK1:O | 3.05555 | Hydrogen Bond |
| N:UNK1:H - A:ASN658:OD1 | 2.40306 | Hydrogen Bond |
| N:UNK1:H - A:ASP690:OD2 | 2.69948 | Hydrogen Bond |
| N:UNK1:H - N:UNK1:O | 1.89695 | Hydrogen Bond |
| A:ARG590:NH1 - N:UNK1 | 3.92092 | Hydrogen Bond; Electrostatic |
| A:ARG590:NH2 - N:UNK1 | 3.64847 | Electrostatic |

\*UNK1: 3,4'-O-Dimethylquercetin; A: Amino acid.

Alphitolic acid (ID_12305768) and arjunolic acid (ID_73641) are the two pentacyclic triterpenoids examined in this study. They generally exhibited fewer binding interactions with the HMG-binding site of HMGR, possibly due to their bulky and less polar triterpene core structure. However, the -OH and carboxylic (-COOH) groups present at both ends of their side chains formed polar hydrogen bond interactions with relevant residues, such as Asp^690^, Lys^691^, Lys^692^, Asn^658^, and Glu^665^. In addition, their pentacyclic rings engaged in non-polar hydrophobic interactions with important residues such as Val^683^, and Lys^691^ (Table 8g-h; supplementary data). Direct studies on the inhibition of HMGR by alphitolic acid and arjunolic acid are currently lacking. However, other triterpenoids, such as oleic acid, eicosyl ester, naringin, apigenin, luteolin, ascorbic acid, and α-tocopherol, have been reported to possess cholesterol-lowering effects (Baskaran *et al*., 2015). Moreover, arjunolic acid has shown protective effects against atorvastatin-induced oxidative stress and apoptosis in renal and hepatic tissues (Pal *et al*., 2015). Its role in activating AMPK and suppressing neuroinflammation in mice with depressive behaviors further suggests it may have an indirect effect on HMGR inhibition (Yang *et al*., 2024).

**Table 8g.** Interaction profile of ID_73641 at HMG-binding site of HMGR.

| HMGR-Phytochemical Interaction | Bond Distance | Category |
| --- | --- | --- |
| A:VAL683 - N:UNK1 | 5.26614 | Hydrophobic |
| N:UNK1:C - A:LYS691 | 3.98626 | Hydrophobic |
| N:UNK1:C - A:LYS691 | 4.19323 | Hydrophobic |
| A:LYS692:HZ3 - N:UNK1:O | 2.08531 | Hydrogen Bond |
| N:UNK1:H - A:ASN658:OD1 | 2.3541 | Hydrogen Bond |
| N:UNK1:C - A:VAL683 | 4.1468 | Hydrophobic |

\*UNK1: Arjunolic acid; A: Amino acid.

**Table 8h.** Interaction profile of ID_12305768 at HMG-binding site of HMGR.

| HMGR-Phytochemical Interaction | Bond Distance | Category |
| --- | --- | --- |
| A:LYS691:HZ1 - N:UNK1:O | 2.89018 | Hydrogen Bond |
| N:UNK1:H - A:ASP690:OD2 | 2.65657 | Hydrogen Bond |
| N:UNK1:H - A:GLU665:OE1 | 2.78814 | Hydrogen Bond |
| N:UNK1:C - A:LYS691 | 3.84679 | Hydrophobic |
| N:UNK1:C - A:VAL683 | 4.42173 | Hydrophobic |

\*UNK1: Alphitollic acid; A: Amino acid.

Cochloxanthin (ID_101202074) is a carotenoid pigment found in certain plants, including Cochlospermum species. It showed polar interactions between its polar side chain (-OH and -COOH groups) and a few HMG-binding residues, such as Asp^690^, Lys^692^, and others Glu^665^. Additionally, hydrophobic bonds were formed between the carbon atoms of its long polyene chain and relevant residues, including Val^683^ and Lys^691^ (Table 8i; supplementary data). These relatively weak binding interactions likely occurred due to the compound’s linear long-chain skeleton, which may not fit properly into the narrow HMG-binding pocket of the enzyme. Metibemu et al. in their in silico study, investigated several carotenoids isolated from *Spondias mombin* and suggested that these compounds possess HMGR inhibitory effects, along with antilipidemic and anticancer properties, but there was no direct link established with cochloxanthin (Metibemu *et al*., 2021). Similarly, Nafiu et al. reported that the polyphenol-rich root extract of *C.planchonii* significantly reduced serum total cholesterol, triglycerides, and LDL levels in hyperlipidemic rats (Nafiu *et al*., 2020). In addition, ethanolic extracts of cochloxanthin and alphitolic acid from the root of *C.tinctorium* have been shown to exhibit antimalarial properties (Ballin *et al*., 2002).

**Table 8i.**
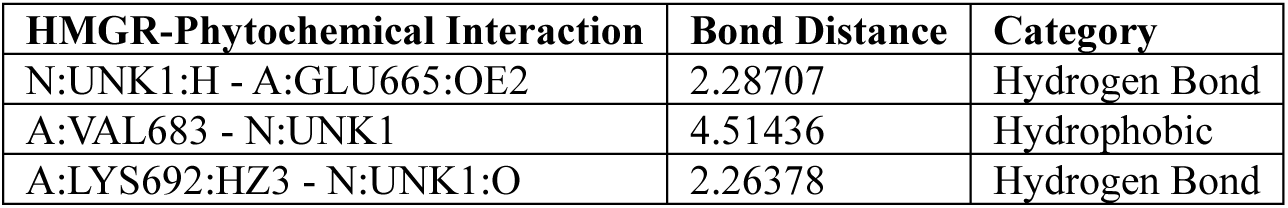
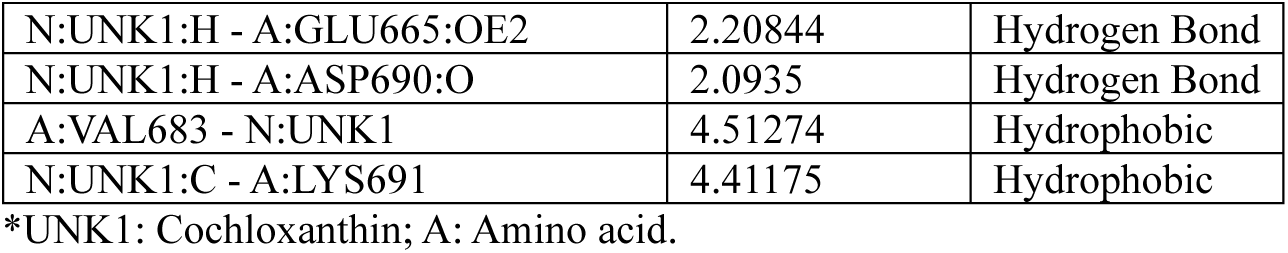
Interaction profile of ID_101202074 at HMG-binding site of HMGR.

3-(Azepan-1-yl)-1,2-benzothiazole 1,1-dioxide (ID_535203) is a heterocyclic sulfonamide derivative with a benzothiazole scaffold and an azepane ring structure, isolated from *C.tinctorium*. Interestingly, this compound revealed promising polar interactions between the sulfonyl functional group (SO2) of its benzothiazole ring and nitrogen atom with HMG-binding residues Arg^590^, Ser^684^, Asp^690^, Lys^692^. Additionally, its benzene and azepane rings formed several catalytically important hydrophobic contacts with residues Val^683^, Lys^691^, Asp^690^, and Met^657^ (Table 8j; Fig. 2D; supplementary data). These interactions suggest the compound may serve as a novel, natural inhibitor of human HMGR. Currently, there is no information available on the effect of this compound on HMGR activities. Nevertheless, a molecular docking study by Ikpa and Tochukwu demonstrated that this compound exhibited higher antiulcer potential than omeprazole by binding strongly to the H^+^/K^+^-ATPase receptor, a key target for proton pump inhibitors. The authors suggested that the compound may have superior gastric proton pump inhibitory potential compared to omeprazole, justifying its traditional use for relieving ulcer in patients (Ikpa and Tochukwu, 2024).

**Table 8j.** Interaction profile of ID_535203 at HMG-binding site of HMGR.

| HMGR-Phytochemical Interaction | Bond Distance | Category |
| --- | --- | --- |
| N:UNK1:N - A:ASP690:OD2 | 3.22982 | Electrostatic |
| A:ARG590:HH11 - N:UNK1:O | 2.1476 | Hydrogen Bond |
| A:SER684:HG - N:UNK1:O | 2.6236 | Hydrogen Bond |
| A:LYS692:HZ2 - N:UNK1:O | 2.36555 | Hydrogen Bond |
| N:UNK1:C - A:ASP690:OD2 | 3.48791 | Hydrogen Bond |
| A:MET657 - N:UNK1 | 5.24197 | Hydrophobic |
| A:LYS691 - N:UNK1 | 4.1143 | Hydrophobic |
| N:UNK1:N - A:ASP690:OD2 | 5.00901 | Electrostatic |
| A:SER684:HG - N:UNK1:O | 1.91383 | Hydrogen Bond |
| A:LYS692:HZ2 - N:UNK1:O | 2.16269 | Hydrogen Bond |
| A:VAL683 - N:UNK1 | 5.41697 | Hydrophobic |
| A:LYS691 - N:UNK1 | 4.89979 | Hydrophobic |
| N:UNK1:N - A:ASP690:OD2 | 5.00901 | Electrostatic |

\*UNK1: 3-(Azepan-1-yl)-1,2-benzothiazole 1,1-dioxide; A: Amino acid.

### Conclusion

This study has identified several bioactive compounds isolated from *C. planchonii* and *C. tinctorium* with potential to inhibit the activity of HMG-CoA reductase (HMGR). The molecular docking results showed that compounds such as ellagic acid and its derivative, flavonoids, triterpenoids, carotenoids, and a benzothiazole derivative, exhibited significant biochemical interactions with the cis-loop residues of the enzyme. This demonstrates the ability of these phytochemicals of interest to serve as natural and safer alternatives for hypercholesterolemia treatment, addressing the limitations posed by synthetic statins.

The findings are also consistent with previous studies that support the cholesterol-lowering and cardioprotective effects of these compounds, either directly or indirectly, through mechanisms such as AMPK activation, HMGR downregulation, and antioxidant properties. While this study provides valuable computational insights into the molecular interactions of these compounds with HMGR, further in vitro and in vitro studies are still necessary to validate their inhibitory potential and therapeutic applications.

## Author Contributions

**Toba I. Olatoye**: Conceptualization, Methodology, Writing-original draft & editing, Writing-review & editing, Software.

## Funding

The author has no funding to declare.

## Data Availability Statement

The raw data file is available to approved researchers and relevant agency upon request.

## Conflict of Interest

The author has no competing interests to declare that are relevant to the content of this article.

## Notes

### Competing Interest Statement

The authors have declared no competing interest.

