## Supplemental figures for "Discovery of Novel Inhibitors of HMG-CoA Reductase using Bioactive Compounds isolated from Cochlospermum Species through Computational Methods"

A

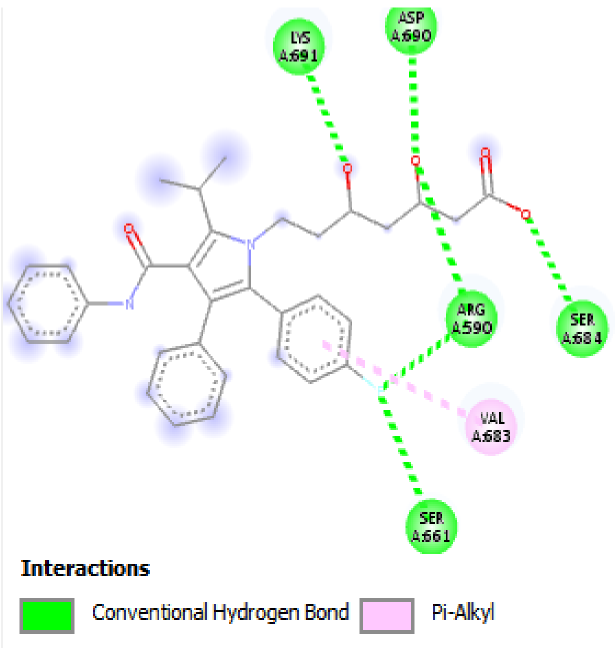
LYS A:691
ASP A:690
ARG A:590
SER A:684
VAL A:683
SER A:661
Interactions
Conventional Hydrogen Bond
Pi-Alkyl

B

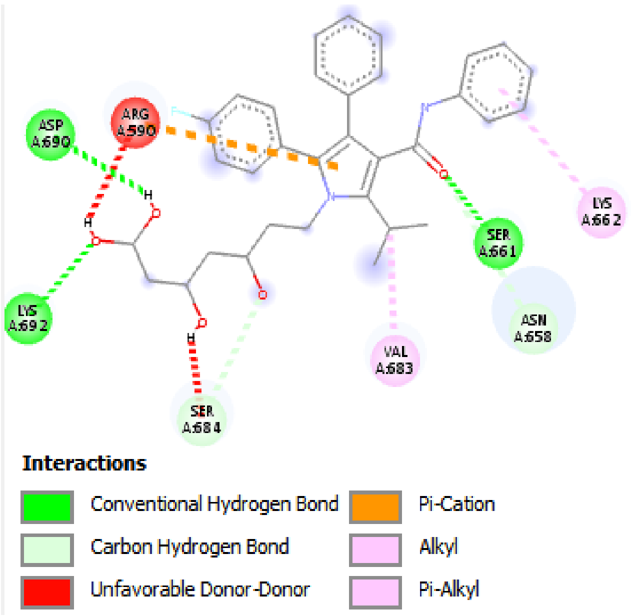
ASP A:690
ARG A:590
LYS A:662
SER A:661
ASN A:658
LYS A:692
VAL A:683
SER A:684
Interactions
Conventional Hydrogen Bond
Carbon Hydrogen Bond
Unfavorable Donor-Donor
Pi-Cation
Alkyl
Pi-Alkyl

C

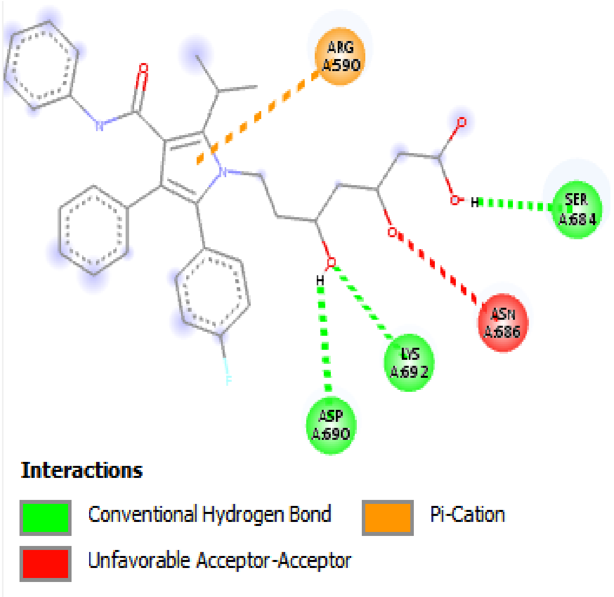
ARG A:590
SER A:684
ASN A:686
LYS A:692
ASP A:690
Interactions
Conventional Hydrogen Bond
Unfavorable Acceptor-Acceptor
Pi-Cation

**D**

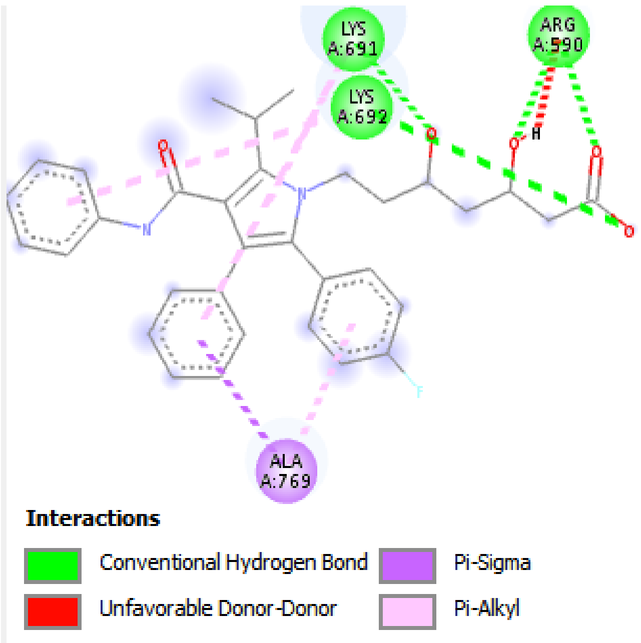
LYS A:691
ARG A:590
LYS A:692
ALA A:769
Interactions
Conventional Hydrogen Bond
Unfavorable Donor-Donor
Pi-Sigma
Pi-Alkyl

**E**

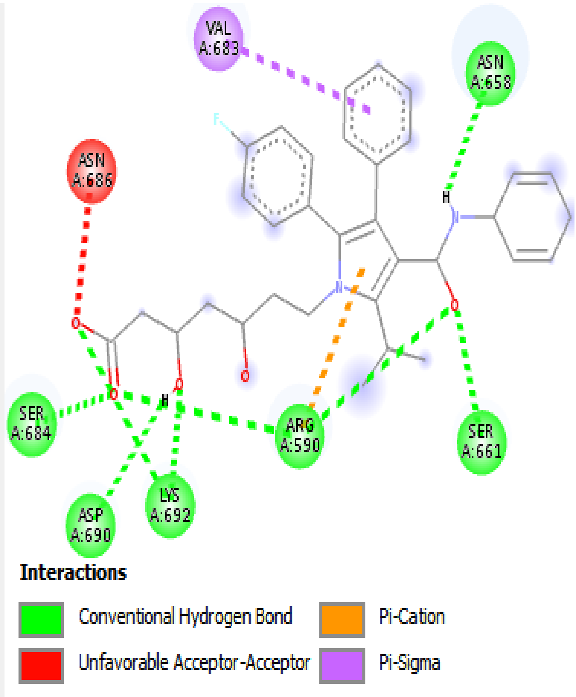
VAL A:683
ASN A:658
ASN A:686
SER A:684
ARG A:590
SER A:661
ASP A:690
LYS A:692
Interactions
Conventional Hydrogen Bond
Unfavorable Acceptor-Acceptor
Pi-Cation
Pi-Sigma

**F**

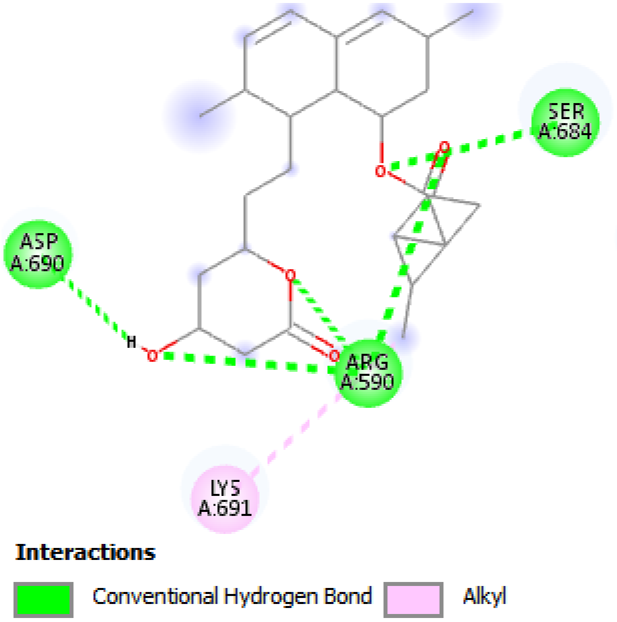
SER A:684
ASP A:690
ARG A:590
LYS A:691
Interactions
Conventional Hydrogen Bond
Alkyl

**G**

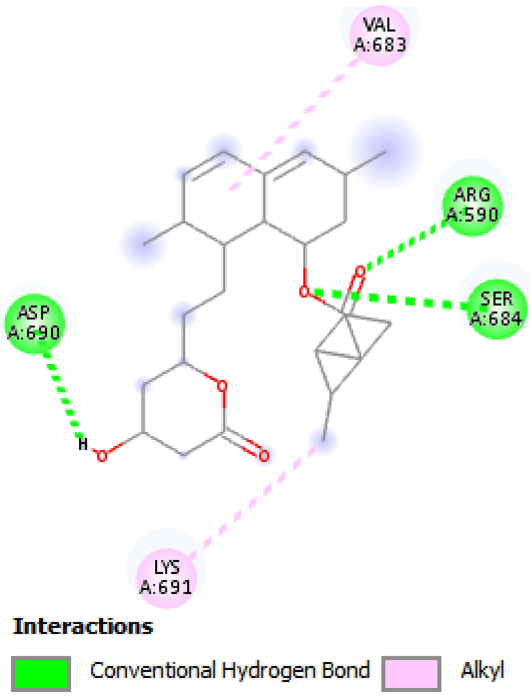
VAL A:683
ARG A:590
SER A:684
ASP A:690
LYS A:691
Interactions
Conventional Hydrogen Bond
Alkyl

H

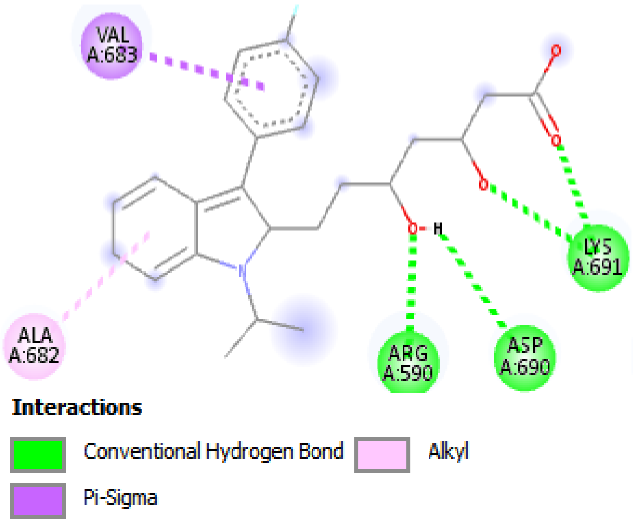
VAL A:683
ALA A:682
ARG A:590
ASP A:690
LYS A:691
Interactions
Conventional Hydrogen Bond
Alkyl
Pi-Sigma

I

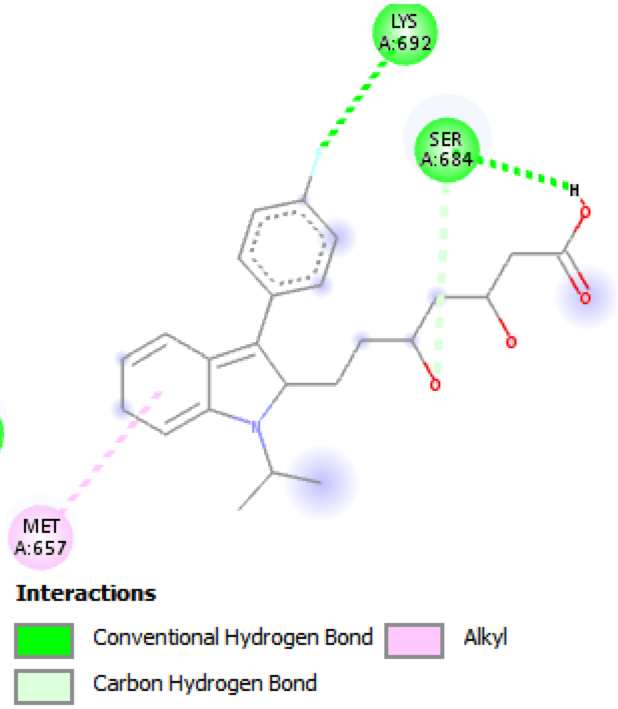
LYS A:692
SER A:684
MET A:657
Interactions
Conventional Hydrogen Bond
Alkyl
Carbon Hydrogen Bond

J

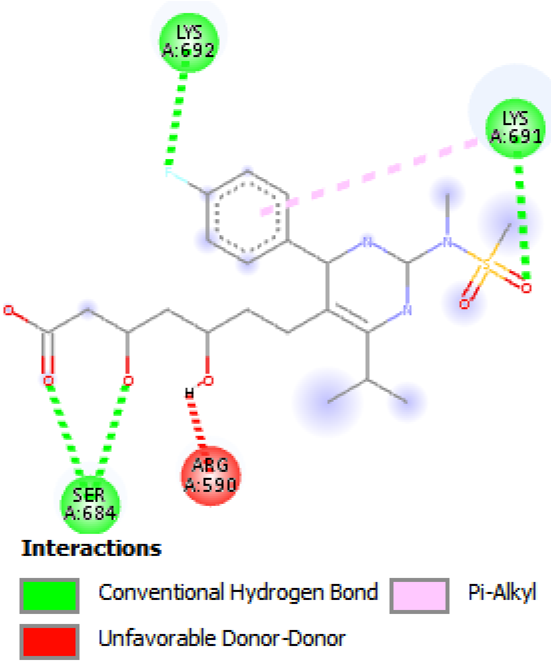
LYS A:692
LYS A:691
SER A:684
ARG A:590
Interactions
Conventional Hydrogen Bond
Pi-Alkyl
Unfavorable Donor-Donor

K

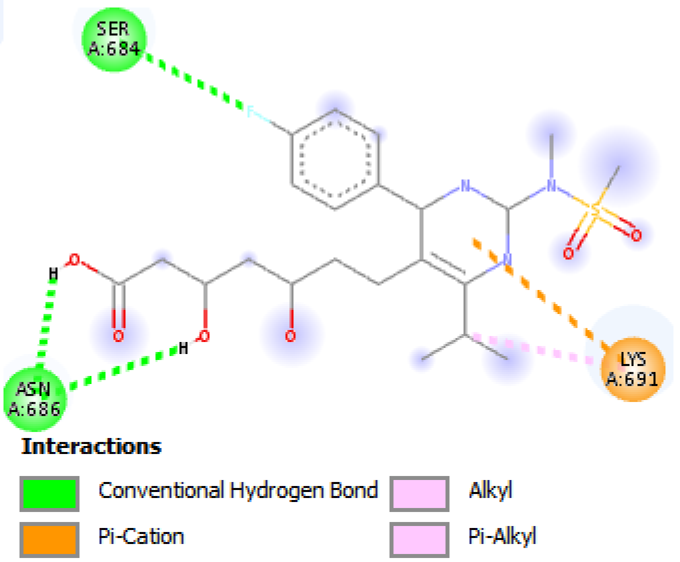
SER A:684
ASN A:686
LYS A:691
Interactions
Conventional Hydrogen Bond
Alkyl
Pi-Cation
Pi-Alkyl

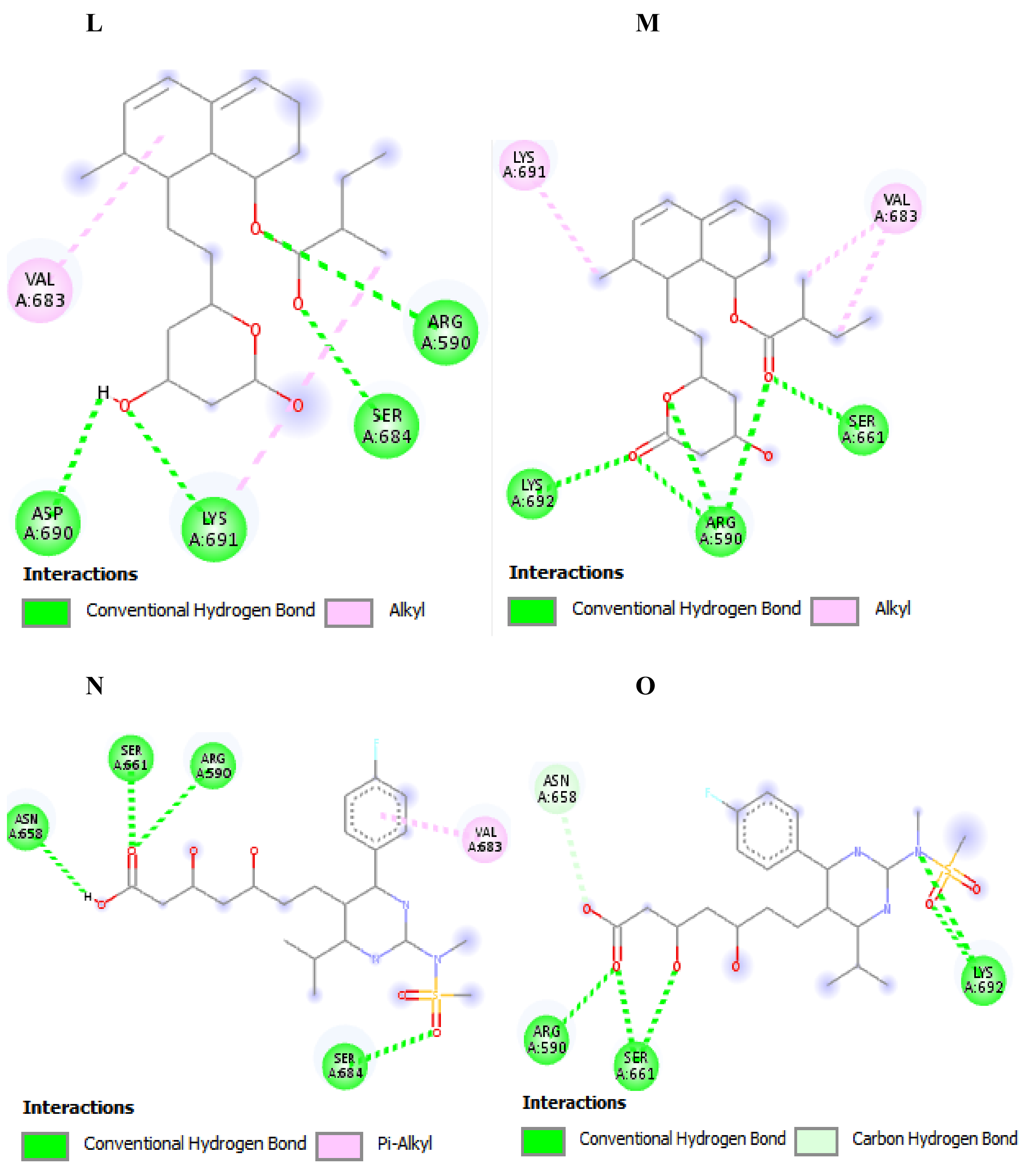


**Supplementary Figure 1:** 2D representation showing the binding interaction of statins with HMG-binding pocket residues of human HMGR. **(A)** 1172 (from literature) **(B)** ID_117/obj01 pose 1 **(C)** ID_117/obj01 pose 2 **(D)** ID_60823 pose 1 **(E)** ID_60823 pose 2 **(F)** ID_54454 pose 1 **(G)** ID_54454 pose 2 **(H)** ID_446155 pose 1 **(I)** ID_446155 pose 2 **(J)** ID_446157 pose 1 **(K)** ID_446157 pose 2 **(L)** ID_64715 pose 1 **(M)** ID_64715 pose 2 **(N)** ID_446156 pose 1 **(O)** 446156 pose 2

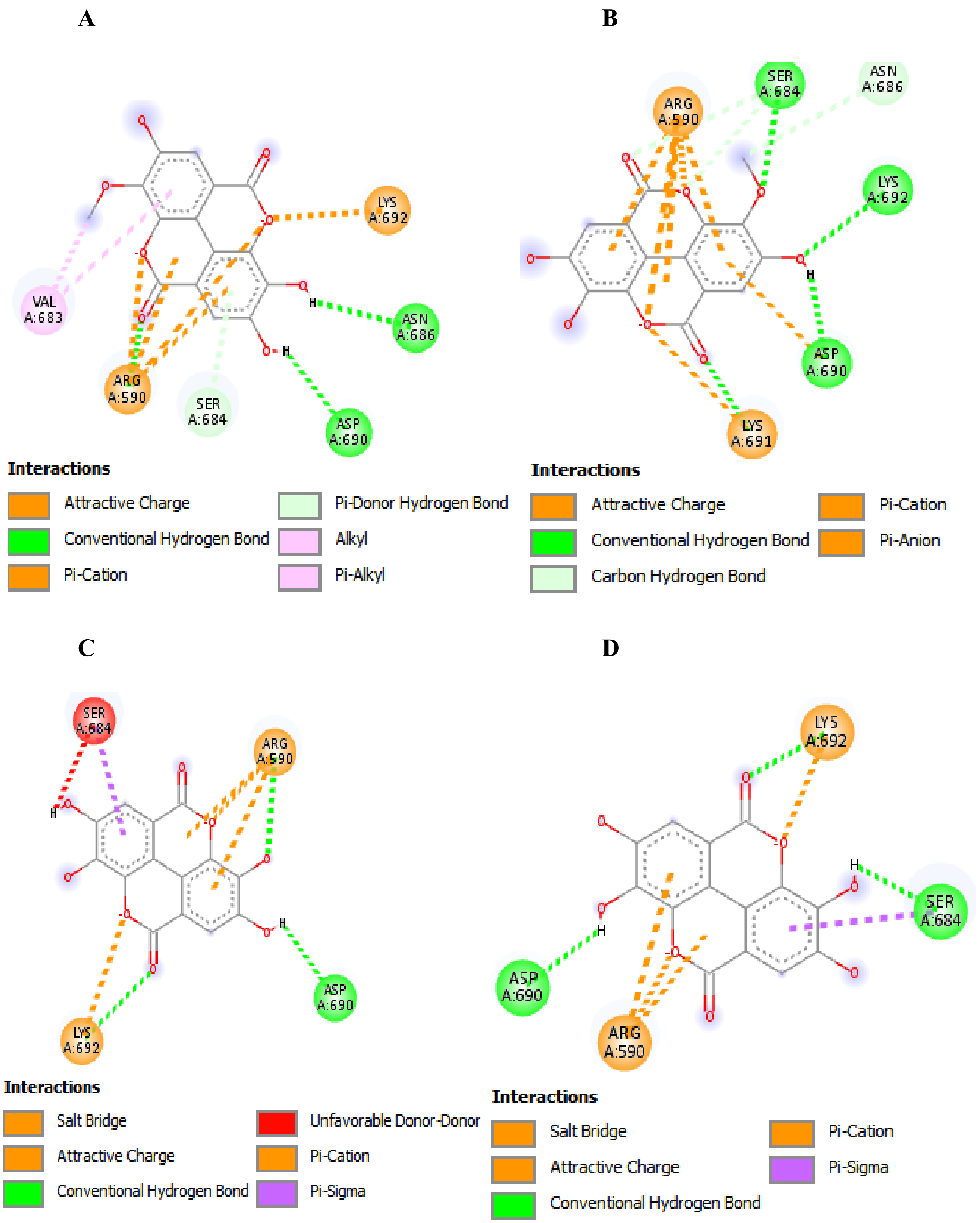


**Supplementary Figure 2:** 2D representation showing the binding interaction of ellagitannins with HMG-binding pocket residues of human HMGR. **(A)** ID_13915428 pose 1 **(B)** ID_13915428 pose 2 **(C)** ID_5281855 pose 1 **(D)** ID_5281855 pose 2

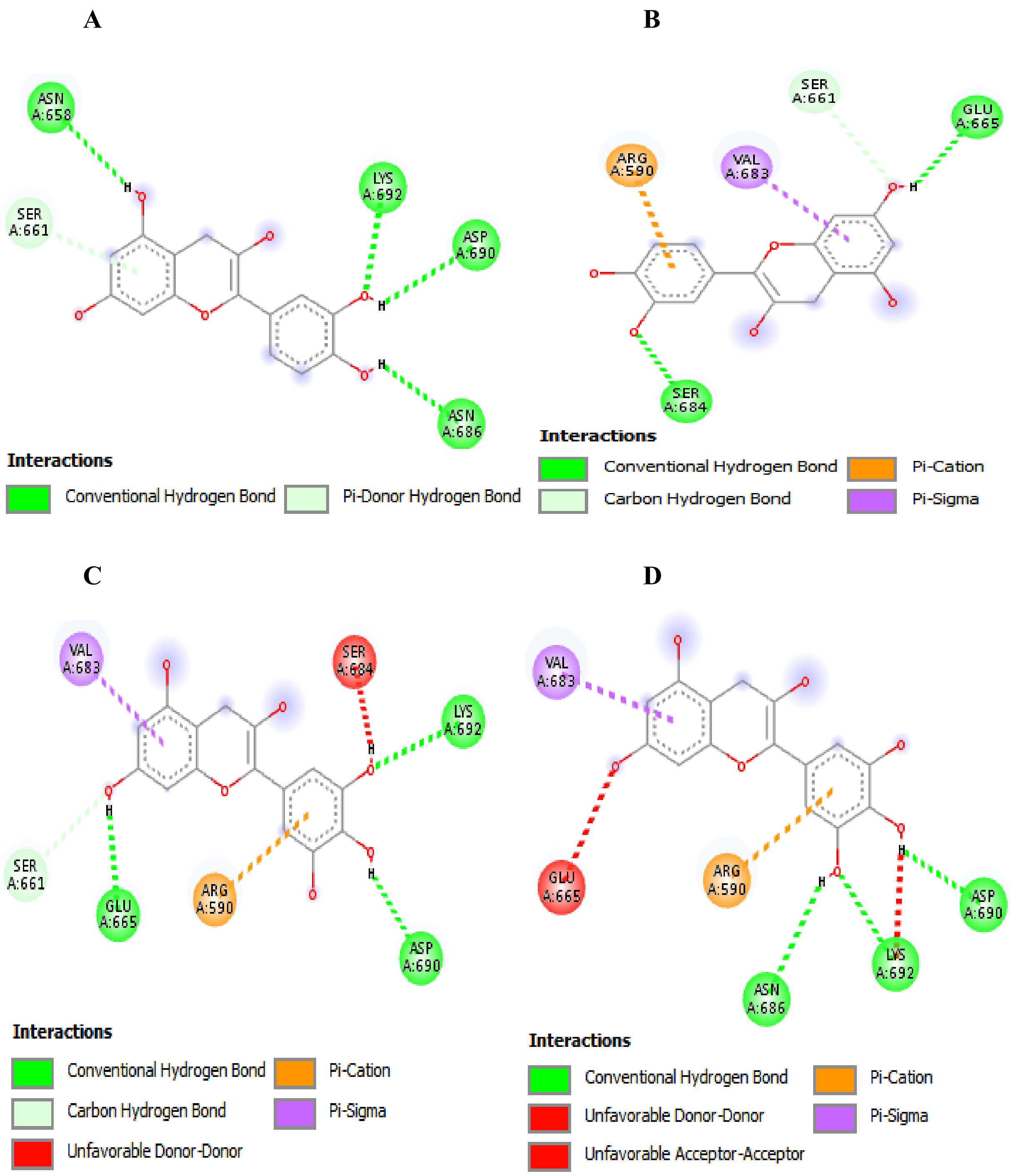
A
ASN A:658
SER A:661
LYS A:692
ASP A:690
ASN A:686
Interactions
Conventional Hydrogen Bond
Pi-Donor Hydrogen Bond
B
SER A:661
GLU A:665
ARG A:590
VAL A:683
SER A:684
Interactions
Conventional Hydrogen Bond
Pi-Cation
Carbon Hydrogen Bond
Pi-Sigma
C
VAL A:683
SER A:684
LYS A:692
SER A:661
GLU A:665
ARG A:590
ASP A:690
Interactions
Conventional Hydrogen Bond
Pi-Cation
Carbon Hydrogen Bond
Pi-Sigma
Unfavorable Donor-Donor
D
VAL A:683
GLU A:665
ARG A:590
ASP A:690
LYS A:692
ASN A:686
Interactions
Conventional Hydrogen Bond
Pi-Cation
Unfavorable Donor-Donor
Pi-Sigma
Unfavorable Acceptor-Acceptor

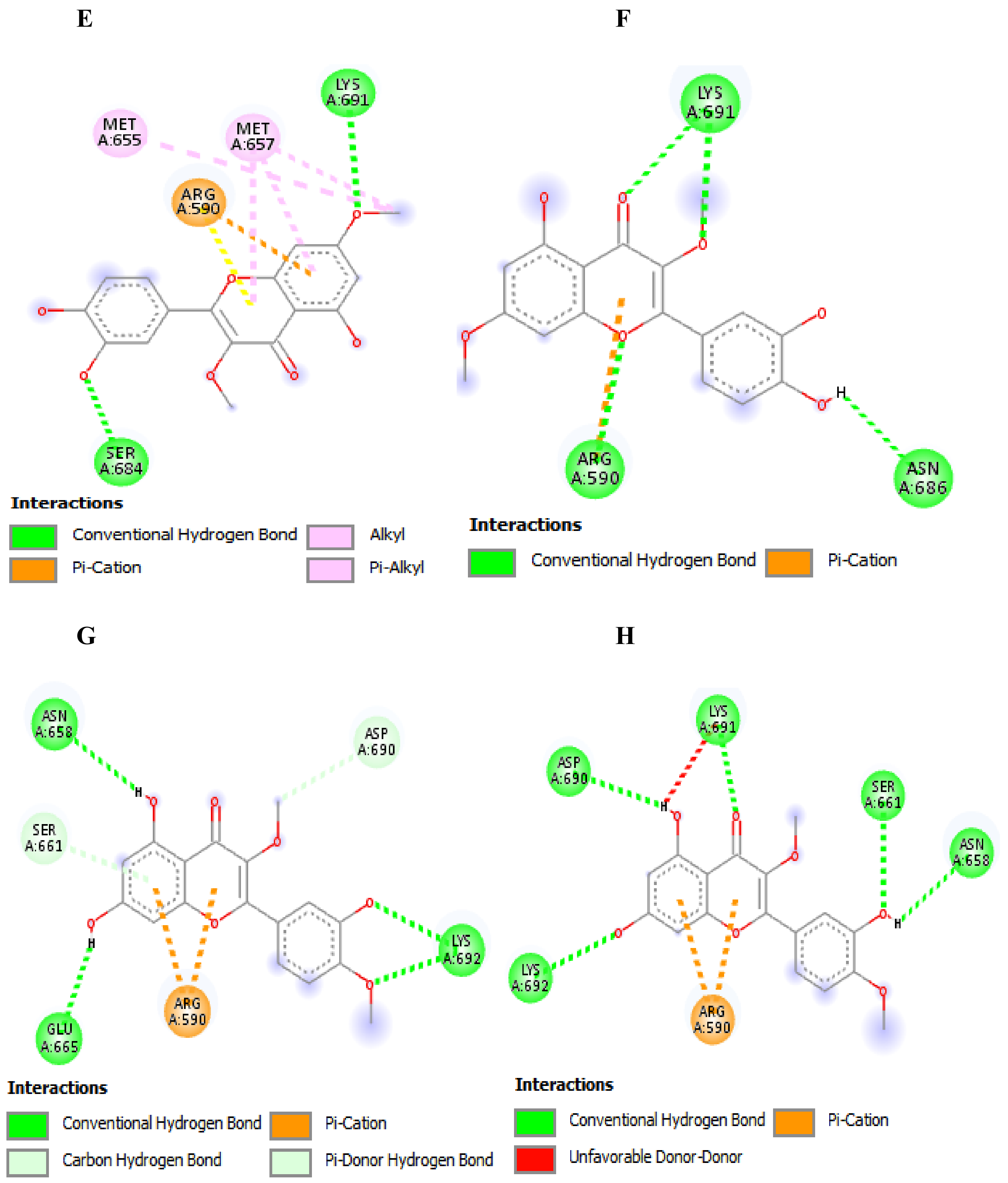


**Supplementary Figure 3:** 2D representation showing the binding interaction of flavonoids with HMG-binding pocket residues of human HMGR. **(A)** ID_9064 pose 1 **(B)** ID_9064 pose 2 **(C)** ID_72277 pose 1 **(D)** ID_72277 pose 2. **(E)** ID_5280417 pose 1 **(F)** ID_5280417 pose 2 **(G)** ID_44446550 pose 1 **(H)** ID_44446550 pose 2

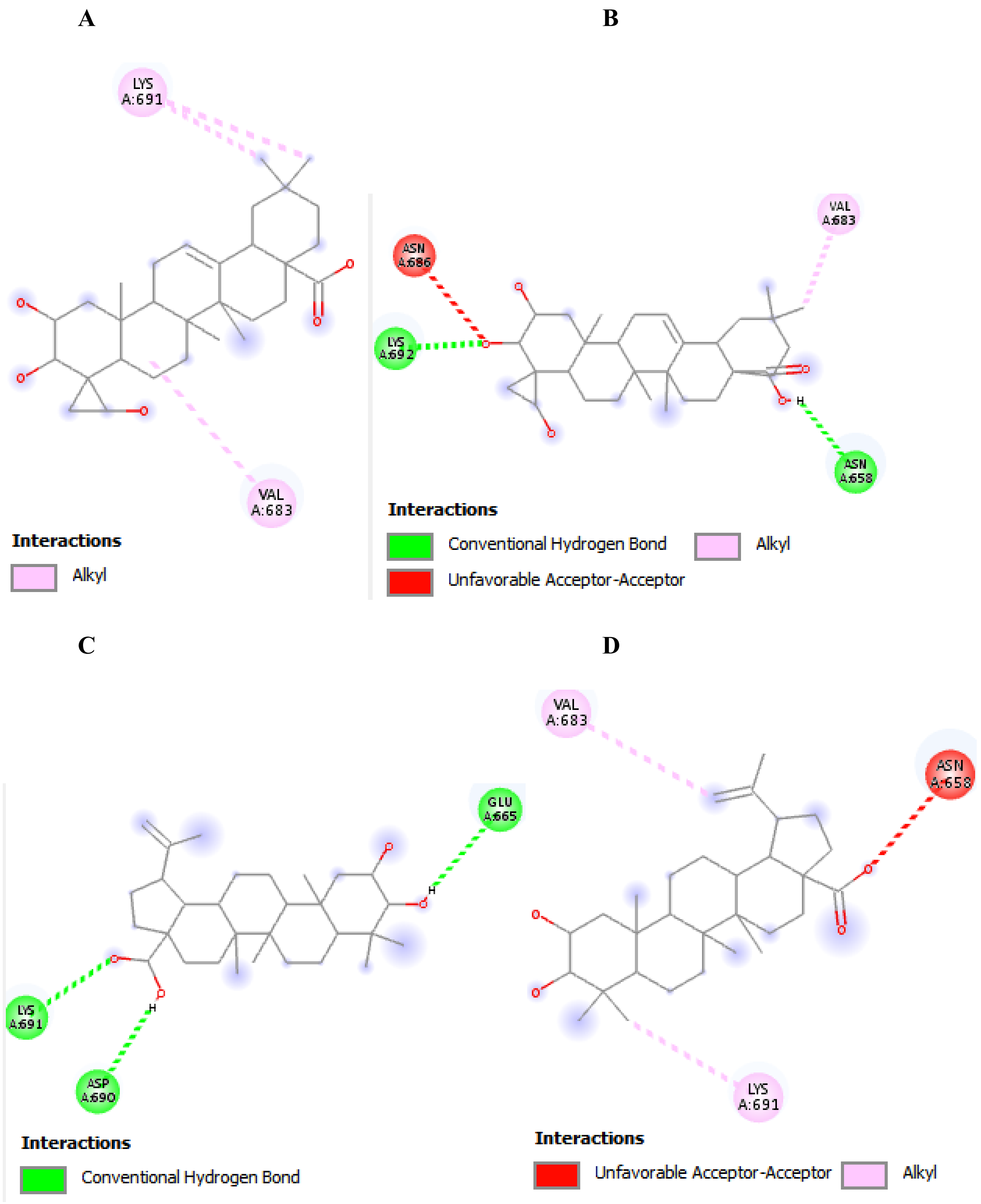


**Supplementary Figure 4:** 2D representation showing the binding interaction of triterpenoid saponins with HMG-binding pocket residues of human HMGR. **(A)** ID_73641 pose 1 **(B)** ID_73641 pose 2 **(C)** ID_12305768 pose 1 **(D)** ID_12305768 pose 2

A B

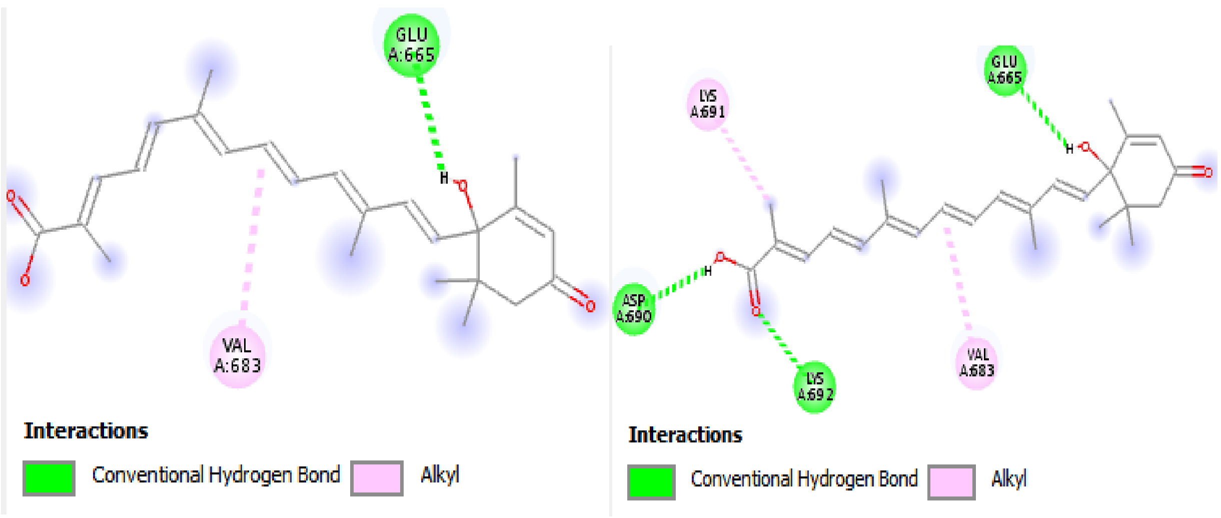


**Supplementary Figure 5:** 2D representation showing the binding interaction of cochloxanthin with HMG-binding pocket residues of human HMGR. **(A)** ID_101202074 pose 1 **(B)** ID_101202074 pose 2

A B

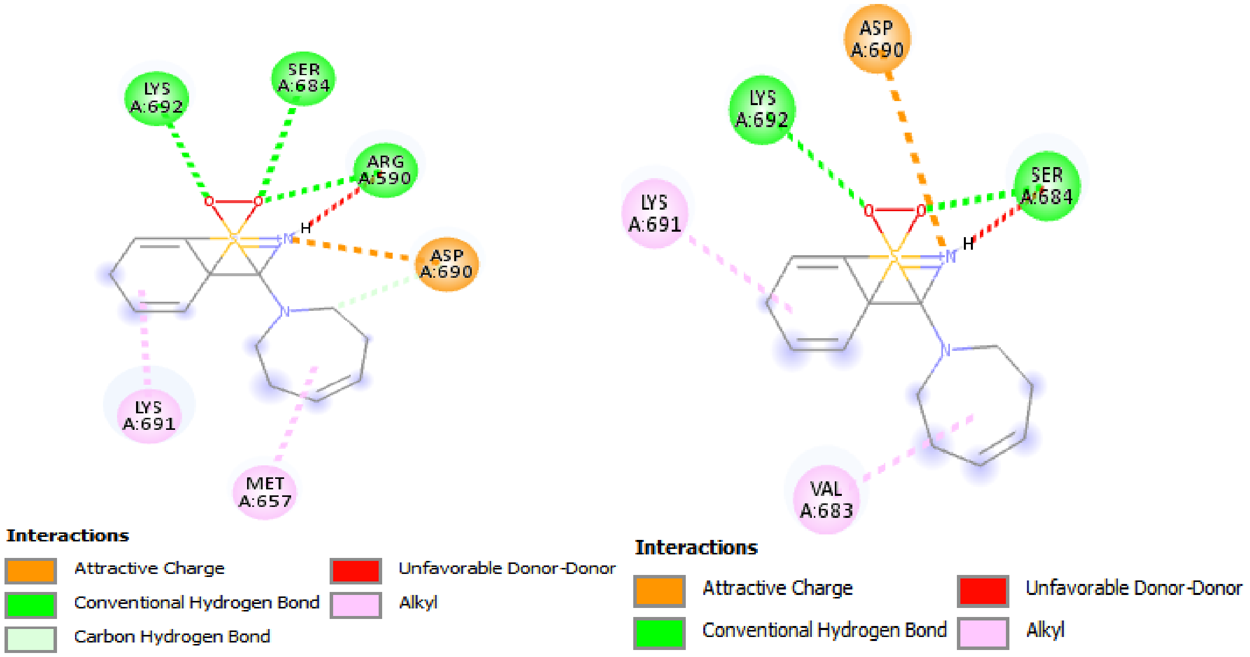


**Supplementary Figure 6:** 2D representation showing the binding interaction of a benzothiazole derivative with HMG-binding pocket residues of human HMGR. **(A)** ID_535203 pose 1 **(B)** ID_535203 pose 2
